# Locality Sensitive Imputation for Single-Cell RNA-Seq Data

**DOI:** 10.1101/291807

**Authors:** Marmar Moussa, Ion I. Măndoiu

**Author notes:** fmarmar.moussa.

## Abstract

One of the most notable challenges in single cell RNA-Seq data analysis is the so called drop-out effect, where only a fraction of the transcriptome of each cell is captured. The random nature of drop-outs, however, makes it possible to consider imputation methods as means of correcting for drop-outs. In this paper we study some existing scRNA-Seq imputation methods and propose a novel iterative imputation approach based on efficiently computing highly similar cells. We then present the results of a comprehensive assessment of existing and proposed methods on real scRNA-Seq datasets with varying per cell sequencing depth.

## 1 Introduction

Emerging single cell RNA sequencing (scRNA-Seq) technologies enable the analysis of transcriptional profiles at single cell resolution, bringing new insights into tissue heterogeneity, cell differentiation, cell type identification and many other applications. The scRNA-Seq technologies, however, suffer from several sources of significant technical and biological noise, that need to be addressed differently than in bulk RNA-Seq.

One of the most notable challenges is the so called drop-out effect. Whether occurring because of inefficient mRNA capture, or naturally due to low number of RNA transcripts and the stochastic nature of gene expression, the result is capturing only a fraction of the transcriptome of each cell and hence data that has a high degree of sparsity. The drop-outs typically do not affect the highly expressed genes but may affect biologically interesting genes expressed at low levels such as transcription factors. Combining cells as a measure to compensate for the drop-out effects could be defeating the purpose of performing single cell RNA-Seq. In this paper we take advantage of the random nature of drop-outs and develop imputation methods for scRNA-Seq. In next section we briefly discuss some existing scRNA-Seq imputation methods and propose a novel iterative imputation approach based on efficiently computing highly similar cells. We then present the results of a comprehensive assessment of the existing and proposed methods on real scRNA-Seq datasets with varying sequencing depth.

## 2 Methods

### 2.1 Existing single cell RNA-Seq imputation methods

#### DrImpute [4]

The DrImpute R package implements imputation for scRNA-Seq based on clustering the data. First DrImpute computes the distance between cells using Spearman and Pearson correlations, then it performs cell clustering based on each distance matrix, followed by imputing zero values multiple times based on the resulting clusters, and finally averaging the imputation results to produce a final value for the drop-outs.

#### scImpute [8]

The scImpute R package makes the assumption that most genes have a bimodal expression pattern that can be described by a mixture model with two components. The first component is a Gamma distribution used to account for the drop-outs, while the second component is a Normal distribution to represent the actual gene expression levels. Thus, in [8], the expression level of gene *i* is considered a random variable with density function *f*_*x*__*i*_ (*x*) = *λ*_*i*_*Gamma*(*x*; *α*_*i*_; *β*_*i*_) + (1 − *λ*_*i*_)*Normal*(*x*; *µ*_*i*_; *σ*_*i*_), where *λ*_*i*_ is the drop-out rate of gene *i*, *α*_*i*_ and *β*_*i*_ are shape and rate parameters of its Gamma distribution component, and *µ*_*i*_ and *σ*_*i*_ are the mean and standard deviation of its Normal distribution component. The parameters in the mixture model are estimated using Expectation-Maximization (EM). The authors’ intuition behind this mixture model is that if a gene has high expression and low variation in the majority of cells, then a zero count is more likely to be a drop-out value than when the opposite occurs, i.e., when a gene has constantly low expression or medium expression with high variation, then a zero count reflects real biological variability. According to [8] this model does not assume an empirical relationship between drop-out rates and mean expression levels and thus allows for more flexibility in model estimation.

#### KNNImpute [16]

Weighted K-nearest neighbors (KNNimpute), a method originally developed for microarray data, selects genes with expression profiles similar to the gene of interest to impute missing values. For instance, consider a gene *A* that has a missing value in cell 1, KNN will find *K* other genes which have a value present in cell 1, with expression most similar to *A* in cells 2 − *N*, where *N* is the total number of cells. A weighted average of values in cell 1 for the *K* genes closest in Euclidean distance is then used as an estimate for the missing value for gene *A*.

There are also some methods for clustering with implicit imputation, like BISCUIT [1,13] and CIDR [9]. These however are out of scope of this paper, as we are focusing on stand-alone imputation methods yielding imputed gene expression profiles that can be used for downstream analyses beyond unsupervised clustering, like dimensionality reduction, counting cells that express known markers, and differential gene expression analysis.

### 2.2 Proposed method: locality sensitive imputation (LSImpute)

We propose a novel algorithm that uses similarity between cells to infer missing values in an iterative approach. The algorithm summary is as follows:

*Step 1.* Given a set *S* of *n* cells (represented by their scRNA-Seq gene expression profiles), start by selecting pairs of cells with highest similarity level until at least *m*_*min*_ distinct cells (*m*_*min*_ = 6 in our implementation) are selected or the highest pair similarity drops below a given threshold. This process guarantees that each selected cell has highest pairwise similarity level to at least one other selected cell.^1^

*Step 2.* Cluster the *m* cells selected in Step 1 using a suitable clustering algorithm (our implementation uses spherical *K*-means with 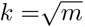). The clusters formed in this step are expected to be “tight”, with each selected cell having high similarity to the other cells in its cluster.

*Step 3.* For each of the clusters identified in step 2, replace zero values for each gene *j* with values imputed based on the expression levels of gene *j* in all the cells within the cluster.

*Step 4.* The selected cells now have imputed values and the clusters they form are collapsed into their respective centroids. The centroids are pooled together with unselected cells to form a new set *S*, and the process is repeated starting again at Step 1.

Note that, naturally, in Step 3 expression levels are imputed only for original cells and not for centroids but centroid expression levels are used in the imputation process if they are selected in Step 1. The expression levels used to replace the zero expression values can be inferred via different models. In Section 3 we give results for two simple approaches, namely using the mean, respectively the median of all expression values for gene *j* in cells belonging to the cluster (these variants are referred to as *LSImputeMean*, respectively *LSImputeMed* in Section 3). Using the median of both zero and non-zero values first, decides implicitly whether a zero is a drop-out event or a true biological effect, and prevents large but isolated expression values from driving imputation of nearby zeros, while collapsing into centroids in each iteration limits the propagation of potential imputation errors. Fig. 1 illustrates the first two steps of the algorithm.

**Fig. 1.**
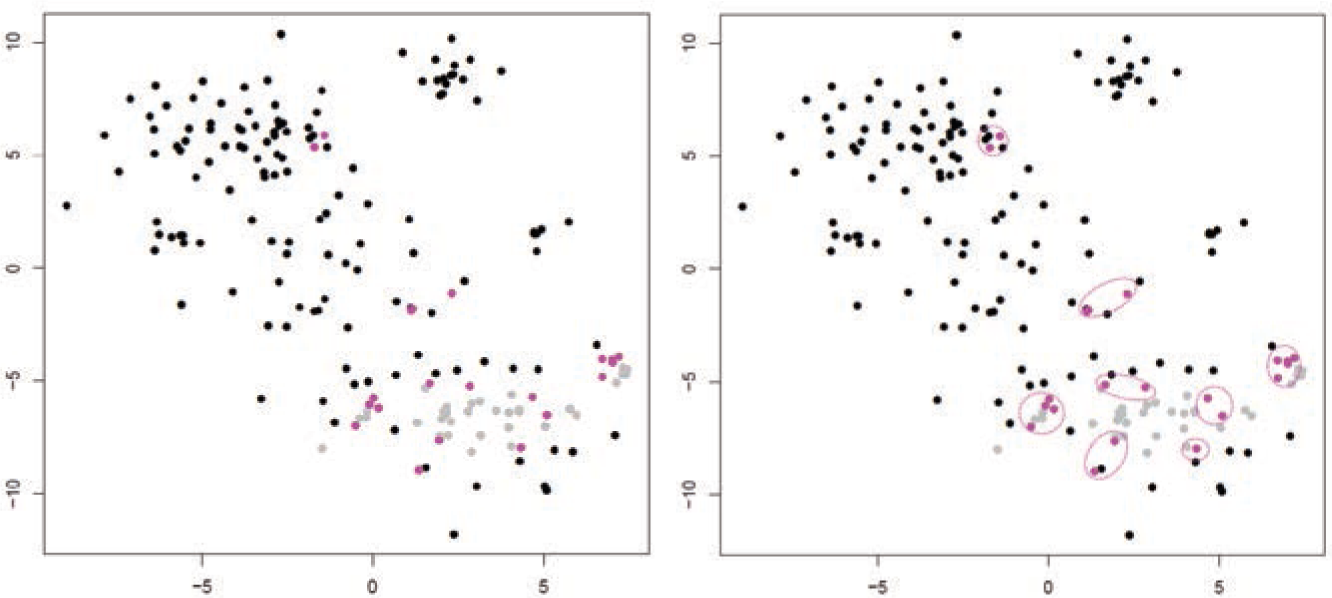
Illustration of Steps 1 (left) and 2 (right) of LSImpute. Gray dots represent already processed cells and collapsed centroids from previous iterations. Pink dots represent cells in pairs with highest similarity level which are selected for clustering.

The worst case number of iterations taken by the algorithm is *O*(*n*) as the total number of remaining cells and centroids starts at *n* and decreases by at least one in each iteration. In practice the number of iterations is much smaller. Our current implementation has two options for finding the pairs of cells with highest similarity level in Step 1. The first option is to use Cosine similarity and the *O*(*n* log *n*) algorithm of [3]. Alternatively, this could be done in *O*(*n*) time using Jaccard similarity and Locality Sensitive Hashing [6]. Both similarity metrics are available in the Shiny app available at http://cnv1.engr.uconn.edu:3838/LSImpute/, where the user can also adjust the minimum similarity threshold used in Step 1. It is recommended however to use a high similarity threshold, which will restrict the imputation to only highly similar cells as a way of being conservative with imputation to avoid the risk of over-imputation. A low similarity threshold can lead to imputing more values and can be used when the data set is of particularly low depths. All results presented in Section 3 use Cosine similarity and a minimum similarity threshold of 0.85 for all sets regardless of depth to avoid over-fitting. Using Jaccard similarity based on the R package LSHR [15] resulted in similar imputation levels as the Cosine similarity based implementation.

### 2.3 Experimental setup

#### Data sets

To assess the performance of the compared imputation methods, we used multiple evaluation metrics on data sets consisting of real scRNA-Seq reads down-sampled to simulate varying sequencing depths per cell. Specifically, we used ultra-deep scRNA-Seq data generated for 209 somatosensory neurons isolated from the mouse dorsal root ganglion (DRG) and described in [7]. An average of 31.5M 2×100 read pairs were sequenced for each cell, leading to the detection of an average of 10,950 ± 1, 218 genes per cell. To simulate varying levels of drop-out effects we down-sampled the full dataset to 50K, 100K, 200K, 300K, 400K, 500K, 1M, 5M, 10M, respectively 20M read pairs per cell. At each sequencing depth *transcript per million (TPM)* gene expression values were estimated for each neuron using the IsoEM2 package [10]. As ground truth we used TPM values determined by running IsoEM2 on the full set of reads. For clustering accuracy evaluation, we used as ground truth the cluster assignment from [7], focusing on the 8 cell populations identified using scRNA-Seq data and not its refinement based on neuron sizes (see Fig. 2). The C1-C8 clusters we use in this paper correspond to the following cell populations identified by their most prominent marker genes as indicated by [7] : C1: Gal; C2: Nppb; C3:Th; C4: Mrgpra3 & Mrgprb4; C5:Mrgprd-high; C6:Mrgprd-low & S100b-high; C7: S100b-low; C8: Ntrk2 & S100b-high.

**Fig. 2.**
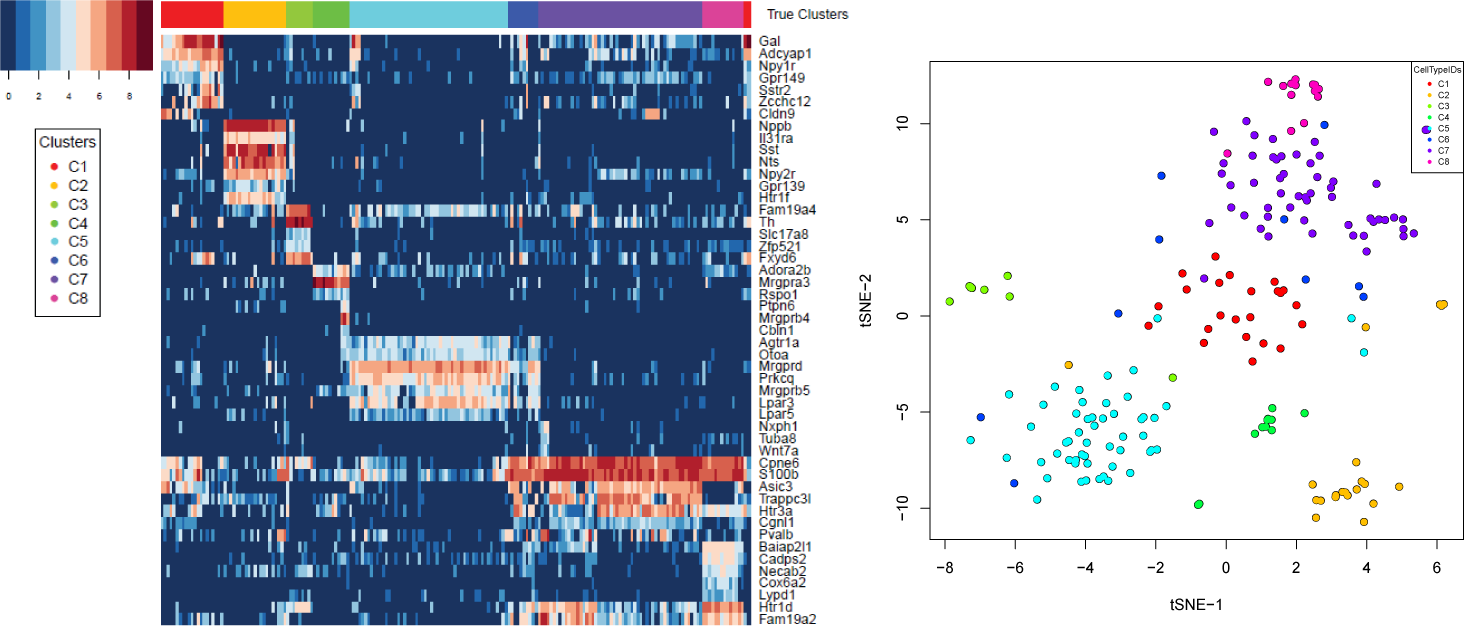
Heatmap of log-transformed TPM values of marker genes identified for DRG neurons in [7] (left) and t-SNE plot showing the 8 clusters from [7] (right).

#### Evaluation metrics

We used the following metrics to evaluate the imputation methods’ performance at different sequencing depths:

- **Detection fraction accuracy**. A common application of single cell analyses is to estimate the percentage of cells expressing a given marker gene, for instance *CD*4+ or *CD*8+ tumor infiltrating lymphocytes [2]. A gene is considered to be detected in a cell if the (imputed or ground truth) TPM is positive. For each imputation method, the detection fraction is defined as the number of cells in which the cell is detected divided by the total number of cells. This was compared to the ‘true’ detection ratio, defined based on ground truth TPM values.
- **Median percent error (MPE)**. As defined in [12], the *Median Percentage Error (MPE)* is the median of the set of relative errors for the gene metric examined, in this case the detection fraction. If a gene has predicted detection fraction *y* and a ground truth detection fraction of *x*, the gene’s relative error is defined as 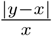. For each sequencing depth we computed MPE relative to all genes as well as subsets of genes corresponding to the four quartiles defined by gene averages of non-zero ground truth TPM values over all cells (ranges of mean non-zero TPM values for the four quantiles were [0, 2.3] (2.3, 6.744], (6.744, 24.517], and (24.517, 18576.98], respectively. Full error curves plotting the percentage of genes with relative error above varying thresholds were also used for a more detailed comparison of imputation methods.
- **Gene detection accuracy**. This metric views gene detection as a binary classification problem. For each imputation method, *true positives (TP)* are the (gene,cell) pairs for which both imputed and ground truth TPM values are positive, while *true negatives (TN)* are (gene,cell) pairs for which both TPM values are zero. The accuracy is computed as the number of true predictions (*TP* + *TN*) divided by the product between the number of genes and the number of cells.
- **Clustering micro-accuracy**. For each sequencing depth and imputation method we clustered imputed TPM values using several clustering algorithms and assessed the effect of imputation on clustering accuracy using the micro-accuracy measure [5,17] defined by 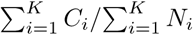, where *K* is the number of classes, *N*_*i*_ is the size of class *i*, and *C*_*i*_ is the number of correctly labeled samples in class *i* relative to the ground truth from [7].

## 3 Results and discussion

To assess imputation accuracy on data sets with varying amounts of drop-outs we sub-sampled the ultra-deep DRG scRNA-Seq data to simulate sequencing depths between 50K and 20M read pairs per cell. For each sequencing depth the metrics described in Section 2.3 were computed for three previous methods (DrImpute, scImpute and KNNImpute), the two variants of our locality sensitive imputation method described in Section 2.2 (LSImputeMean and LSImputeMed), and, as a reference, for the ‘Raw Data’ consisting of TPM values without any imputation.

### Detection fraction accuracy

Fig. 3 plots the true detection fraction (*x*-axis) against the detection fraction in the raw data, respectively after imputation with each of the five compared methods (*y*-axis) at three selected sequencing depths (100K, 1M, respectively 10M read pairs per cell; high resolution plots for all ten evaluated sequencing depths are available in Appendix A. Each dot in the scatter plots represents one gene. Dot color shades are based on the four quartiles as defined above. For an ideal imputation method all dots would lie on the main diagonal, which represents perfect agreement between predicted and true detection fractions. Dots below the diagonal correspond to genes for which the detection fraction is under-estimated, while dots above the diagonal correspond to genes for which the detection fraction is over-estimated. Dropouts in the raw data yield severe under-estimation of the detection fraction for most genes at sequencing depths of 100K and 1M read pairs per cell, but at 10M read pairs per cell detection fractions computed based on raw data are very close to the true fractions for nearly all genes. Existing methods over-impute detection fractions for most genes, even at low sequencing depths. At 100K read pairs per cell LSImputeMed under-estimates detection fractions, improving very little over raw values, while LSImputeMean gives most accurate detection fractions. At higher sequencing depths LSImputeMean begins over-imputing, while LSImputeMed yields most accurate detection fractions at 1M read pairs per cell and only slightly over-imputes at 10M read pairs per cell.

**Fig. 3.**
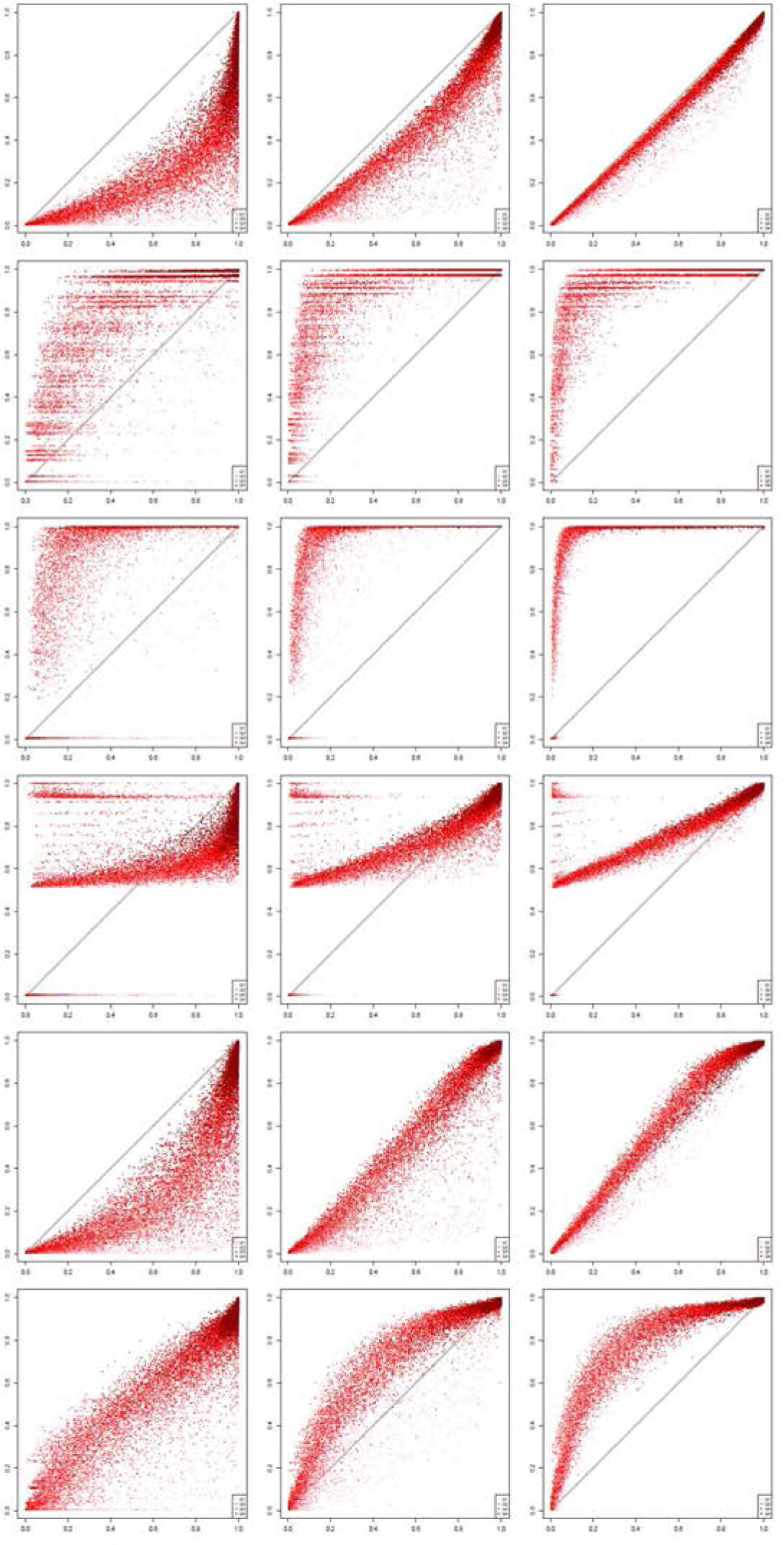
True vs. imputed detection fractions (left to right: 100K, 1M, 10M read pairs per cell; top to bottom: Raw Data, DrImpute, scImpute, KNNImpute, LSImputeMed, and LSImputeMean).

### Detection fraction error curves and MPE comparison

While dot-plots in Fig. 3 give a useful qualitative comparison of detection fraction accuracy of different methods, for a more quantitative comparison of detection fraction accuracy Fig. 4 gives the so called *error curve* of each method. The error curve plots, for every threshold *x* between 0 and 1, the percentage of genes with a relative error above *x*. The error curves in Fig. 4 confirm that LSImputeMean has highest detection fraction accuracy of the compared methods at a sequencing depth of 100K read pairs per cell, while LSImputeMed significantly outperforms the other methods at 1M read pairs per cell and matches raw data accuracy at 10M read pairs per cell. The relative performance of the methods can be even more concisely captured by their MPE values, which are the abscissae of the points where the horizontal line with an ordinate of 0.5 crosses the corresponding error curves. The surface plots in Fig. 5 display MPE values (*y*-axis, on a logarithmic scale) as a function of both sequencing depth (*x*-axis) and mean non-zero expression quartile (*z*-axis). The only imputation methods that do not result in MPE values over 100%, depicted in red in the surface plot, are LSImputeMed and LSImputeMean. At all sequencing depths and for all assessed imputation methods genes in the lowest quartile (Q1) have very high MPE, suggesting that detection fractions based on imputed values should not be used for these genes.

**Fig. 4.**
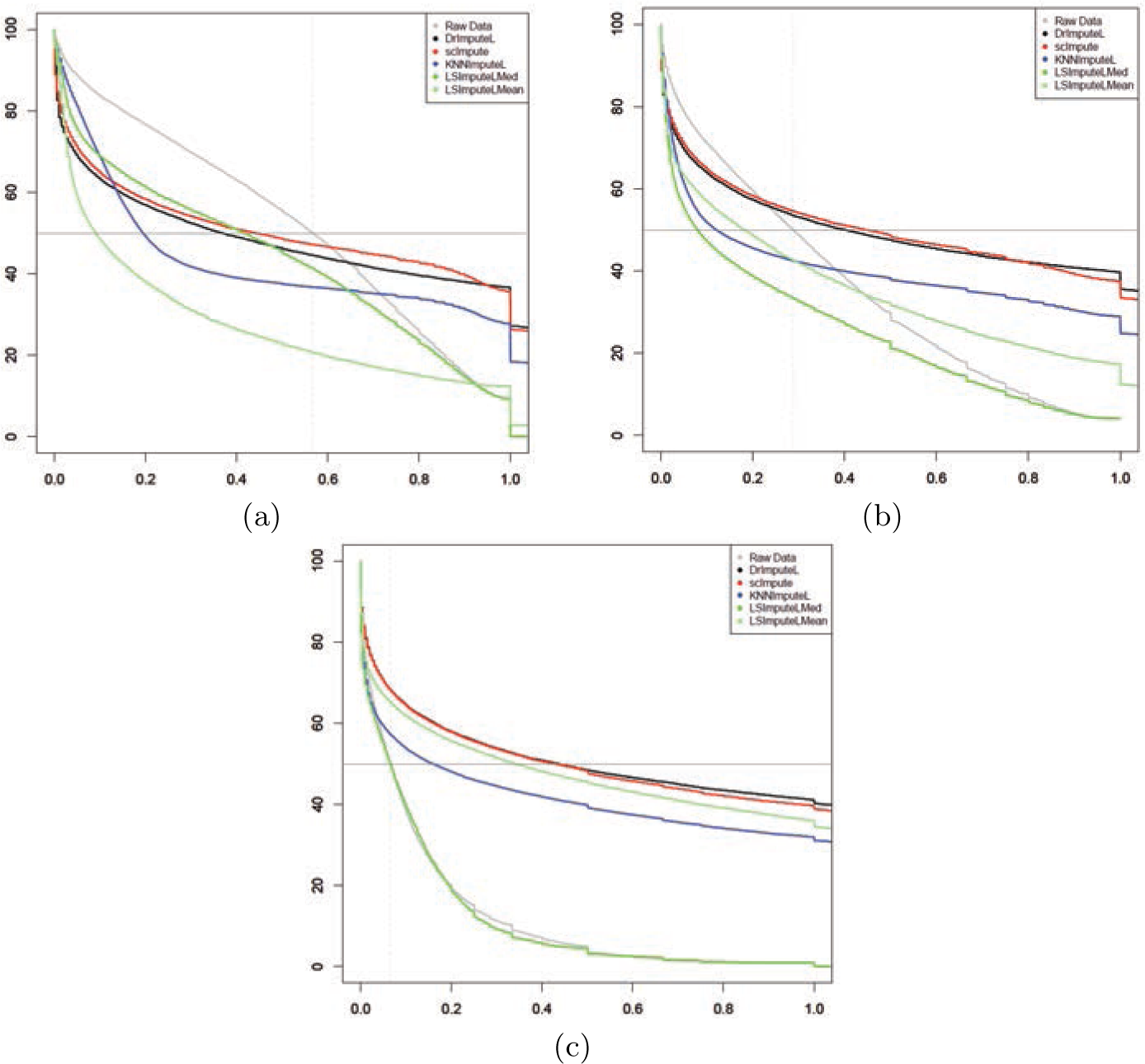
Error curves for (a) 100K, (b) 1M, respectively (c) 10M read pairs per cell. The abscissa of dashed vertical lines correspond to MPE of raw data.

**Fig. 5.**
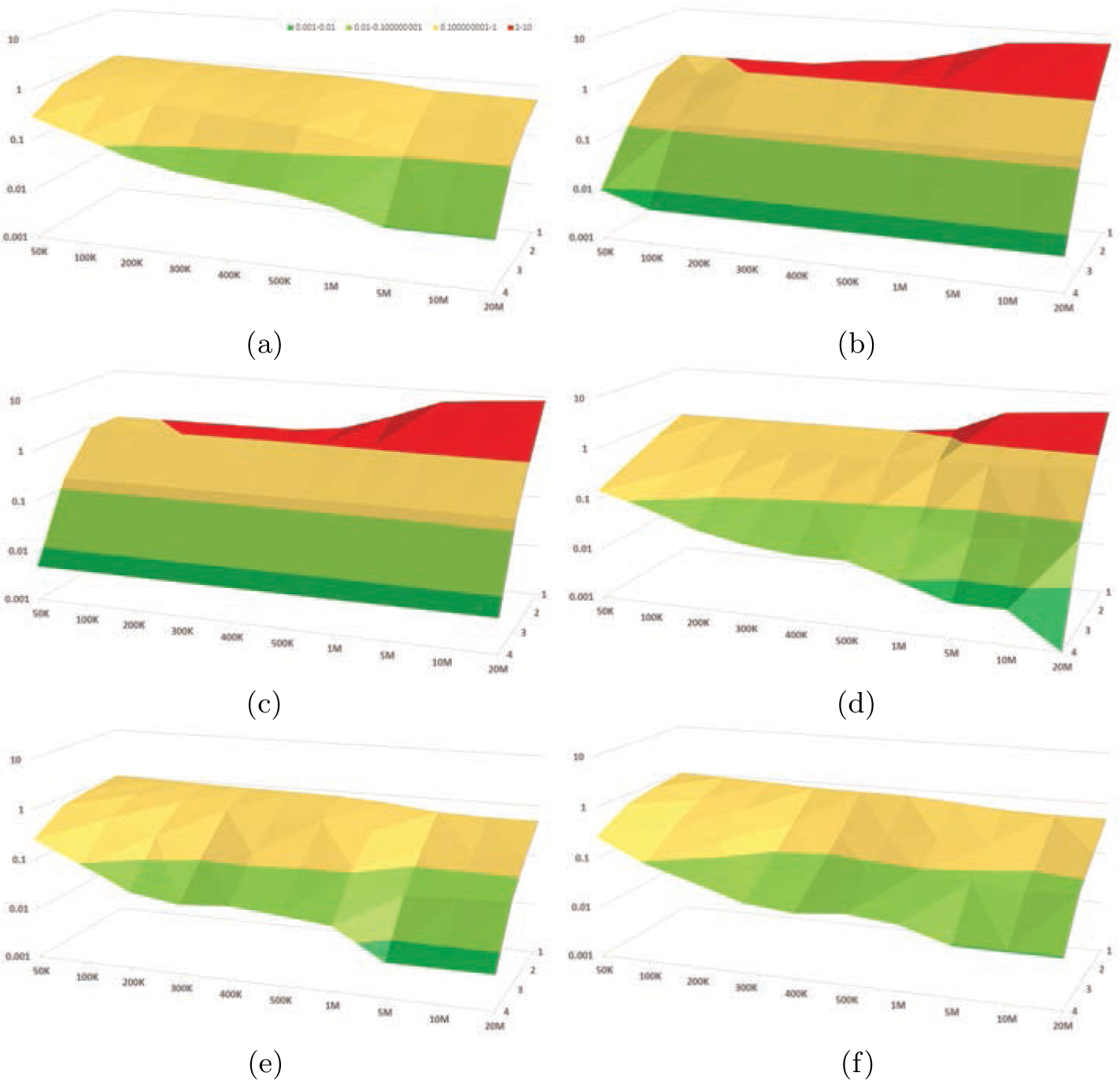
Surface plots indicating Median Percent Error values in log scale (y-axis) for each depth (x-axis) in each quantile (z-axis) for each method: (a) Raw data, (b) DrImpute, (c) scImpute, (d) KNNImpute, (e) LSImputeMed, and (f) LSImputeMean

### Gene detection accuracy and relation to MPE

Table 1 shows the gene detection accuracy achieved by the compared imputation methods, with the highest accuracy at each sequencing depth typeset in bold. We assessed gene detection accuracy both based on fractional ground truth and imputed TPM values, as well as after rounding both to the nearest integer, which is equivalent to using a TPM of 0.5 as the detection threshold. For the results without rounding, DrImpute has the highest gene detection accuracy at 50K and 100K read pairs per cell. LSImputeMean has highest gene detection accuracy for 200K read pairs per cell, while LSImputeMed outperforms the other methods for 300K-1M read pairs per cell. Raw data (no imputation) gives best gene detection accuracy at 5M read pairs per cell and higher depths. For the rounded data sets, DrImpute also has the highest gene detection accuracy at 50K and 100K read pairs per cell, while LSImputeMed outperforms the other methods for 200K-500K read pairs per cell. For sequencing depth of 1M read pairs per cell and higher the raw data gives best detection accuracy followed by LSImpute methods.

**Table 1.**
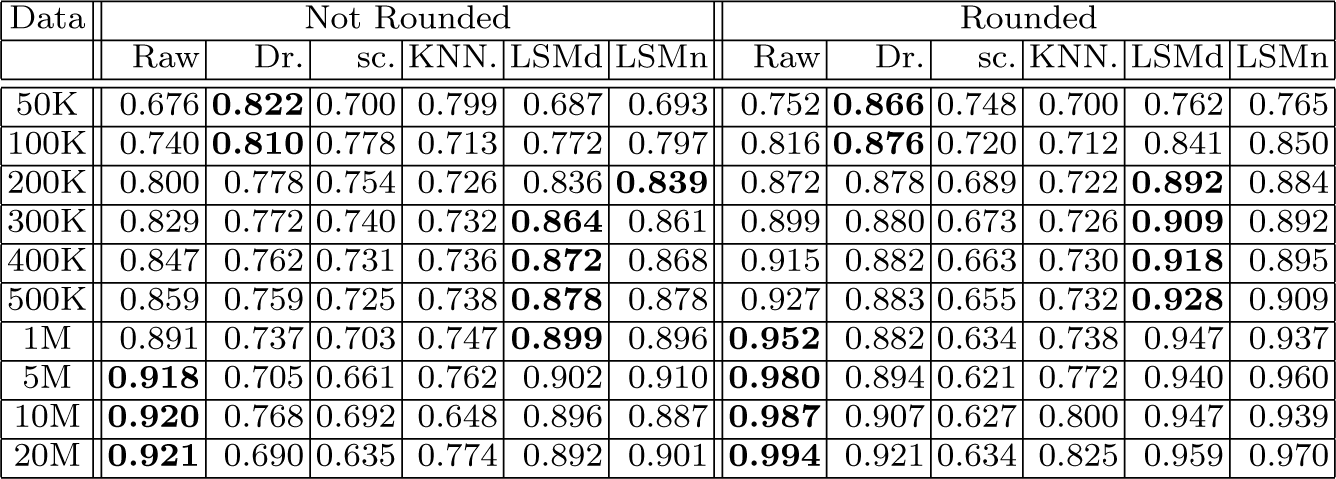
Gene detection accuracy

At very low sequencing depth it is possible for some methods to impute values that are not detected in the ground truth. This could lead to good performance in detection fraction accuracy despite low performance in gene detection accuracy. Furthermore, although one would expect all accuracy measures to improve with increased sequencing depth, this may not necessarily be the case for methods that over-impute. To illustrate the relation between MPE and gene detection accuracy and the effect of sequencing depth increase, in Fig. 6 we plot for each method the gene detection accuracy and MPE achieved without rounding at each sequencing depth from 50K up to 20M read pairs per cell, with consecutive depths connected by arrows pointing in the direction of sequencing depth increase. Since high accuracy and low MPE are preferable, the points near the lower right corner of the plot and arrows pointing towards it indicate better results. For some methods like scImpute and DrImpute, although the starting point (50K read pairs per cell) shows considerable improvement over raw data, as sequencing depth increases one or both of the accuracy measures substantially worsen due to over-imputation. Both LSImputeMed and LSImputeMean start with improvement over raw data in both MPE as and Gene Detection Accuracy and continue in the right direction for higher depths until as mentioned before, the raw data without any imputation gives slightly better gene detection accuracy at 5M read pairs per cell and higher, which suggests that imputation at such high depths comes with the risk of over-imputation for all methods tested.

**Fig. 6.**
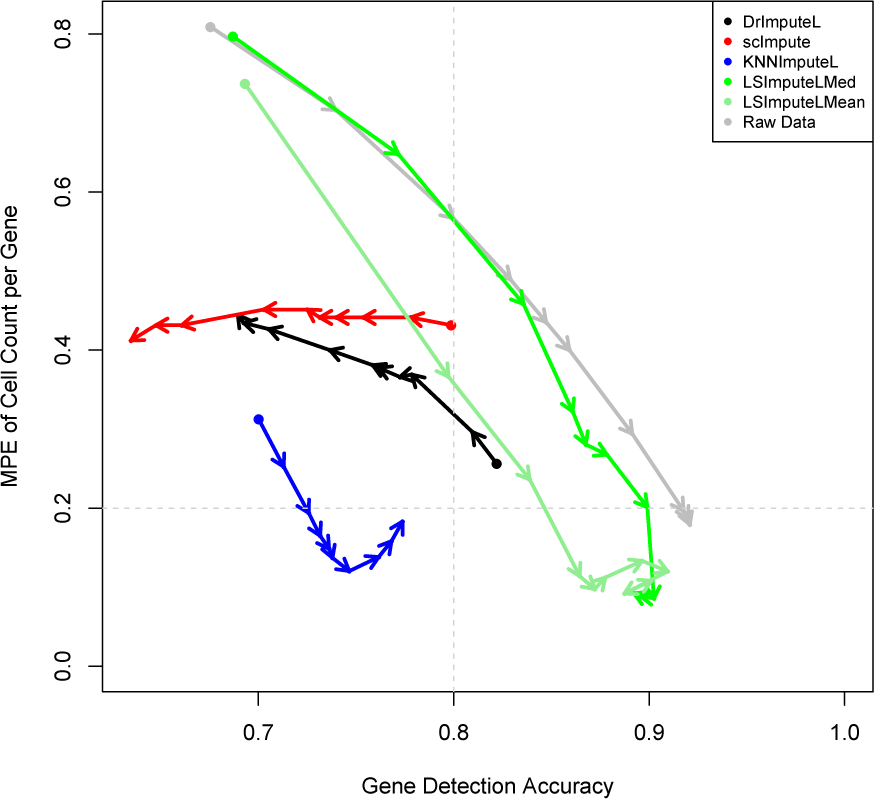
Gene detection accuracy vs. MPE at varying sequencing depths.

### Clustering accuracy

In order to assess the impact of imputation on clustering results, we tested each of the imputation methods in combination with following clustering methods: PCA based hierarchical clustering using Spearman correlation, the TF-IDF Top C clustering approach from [11], and PCA based spherical *k*-means clustering. The micro-accuracy results in Fig. 7 suggest that the effect of imputation varies when combined with different clustering approaches. We also tested Seurat [14] *k*-means clustering of genes and cells (using *k* = 8 with default parameters), however there was very little change in clustering accuracy for different depths. Although the MPE and detection accuracy of some imputation methods suggest the imputation radically alters gene expression profiles, the similarity between cells of a cluster could still hold when all cell profiles are changed in a consistent manner. This can very well lead to no or little change in clustering accuracy, when in fact cell expression profiles are far from the ground truth as the MPE and gene detection accuracy results suggest. As seen in Fig. 8 featuring the log(*x* + 1) expression levels of the marker genes for the DRG 100K data set, although the expression levels of most genes are changed through imputation, the clusters driven by high expression levels of several marker genes can still be the prominent signal for clustering and in most cases this signal remains visually apparent in the heatmaps. Clustering accuracy is hence not recommended as the sole performance evaluation metric when assessing imputation methods.

**Fig. 7.**
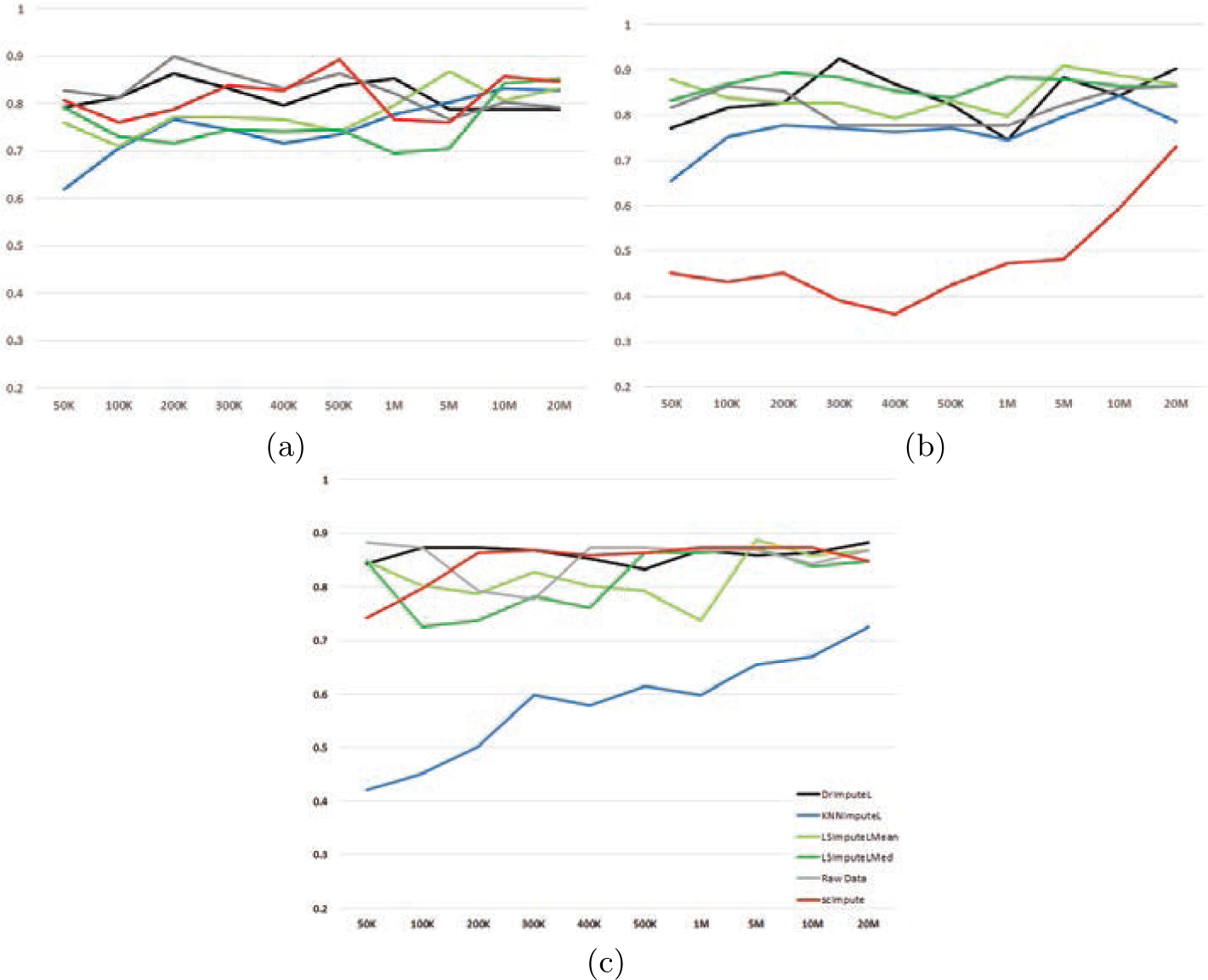
Micro-accuracy on inputed data for (a) PCA-based hierarchical clustering using Spearman correlation, (b) TF-IDF_Top_C [11], and (c) PCA-based spherical k-means.

**Fig. 8.**
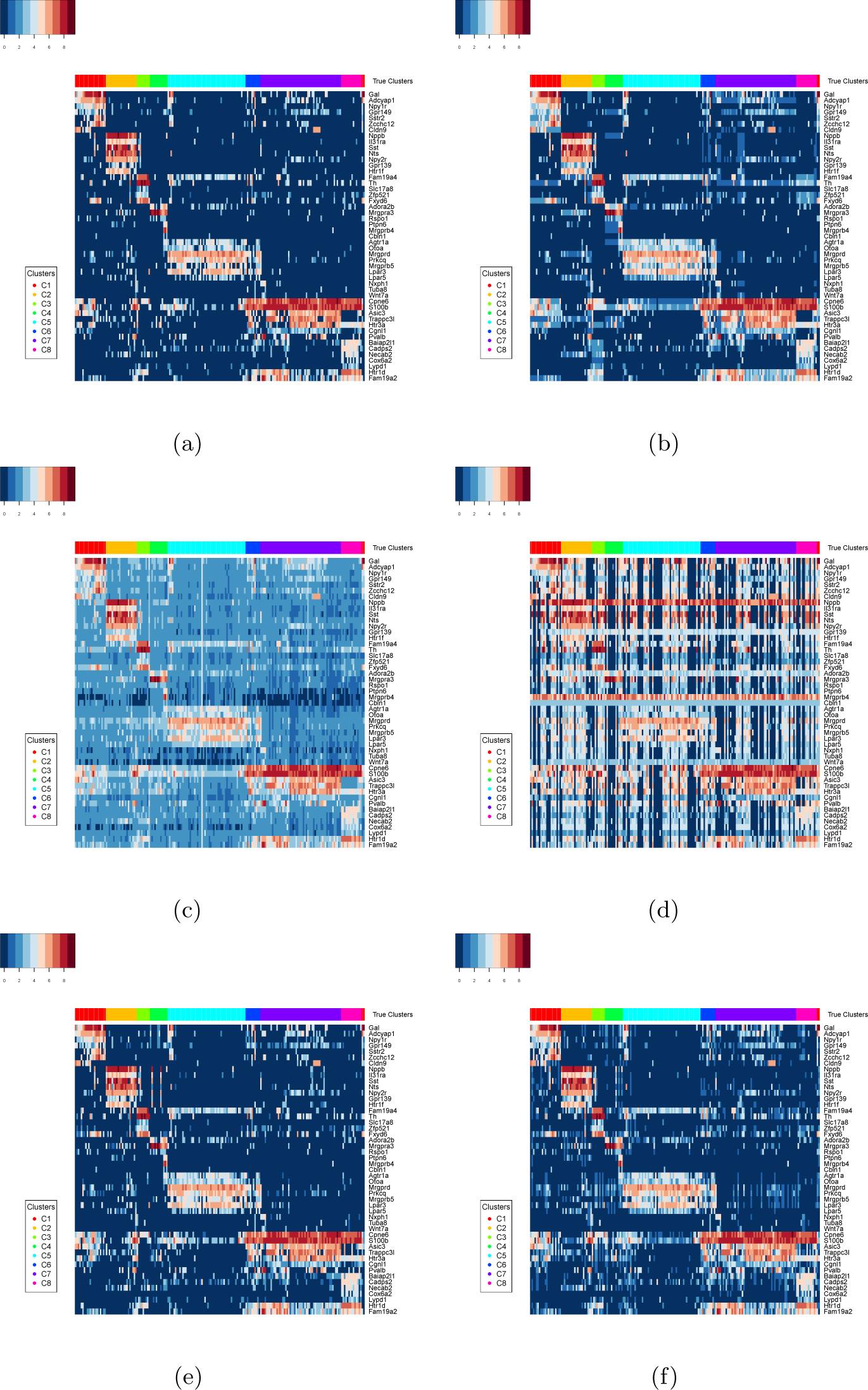
Heatmaps of marker genes from [7] for the 100K DRG dataset: (a) Raw Data, (b) DrImpute, (c) scImpute, (d) KNNImpute, (e) LSImputeMed, (f) LSImputeMean.

## 4 Conclusion

Although imputation can be a useful step in scRNA-Seq analysis pipelines, it can become a two-edged sword if expression values are over-imputed. In this paper we evaluated the performance of several existing imputation R packages and presented a novel approach for imputation. LSImpute, especially the variant based on median imputation, tends to impute more conservatively than existing methods resulting in improved performance based on a variety of metrics. Overall, LSImpute is more likely to reduce drop-out effects and reduce sparsity of the data without introducing false expression patterns or over-imputation. Cosine and Jaccard similarity based implementations of LSImpute are available as a Shiny app at http://cnv1.engr.uconn.edu:3838/LSImpute/.

## Acknowledgements

This work was partially supported by NSF Award 1564936, NIH grants 1R01MH112739-01 and 2R01NS073425-06A1, and a UConn Academic Vision Program Grant. IIM is a co-founder and holds an interest in SmplBio LLC, a company developing cloud-based scRNA-Seq analysis software. No products, services, or technologies of SmplBio have been evaluated or tested in this work.

## A High resolution plots of true vs. imputed detection fractions for all ten evaluated sequencing depths)

**Fig. 9.**
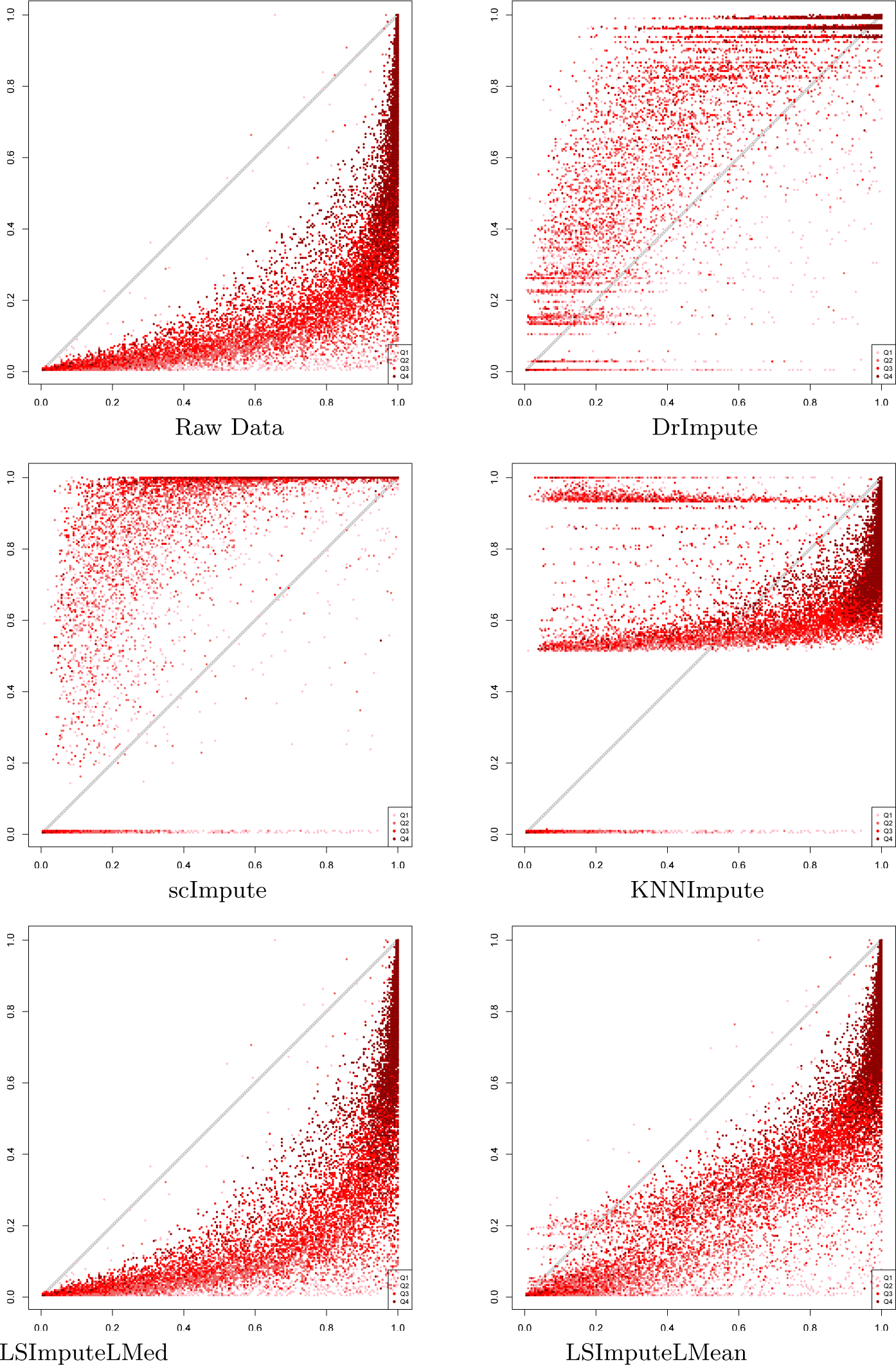
True vs. imputed detection fractions for 50K read pairs per cell.

**Fig. 10.**
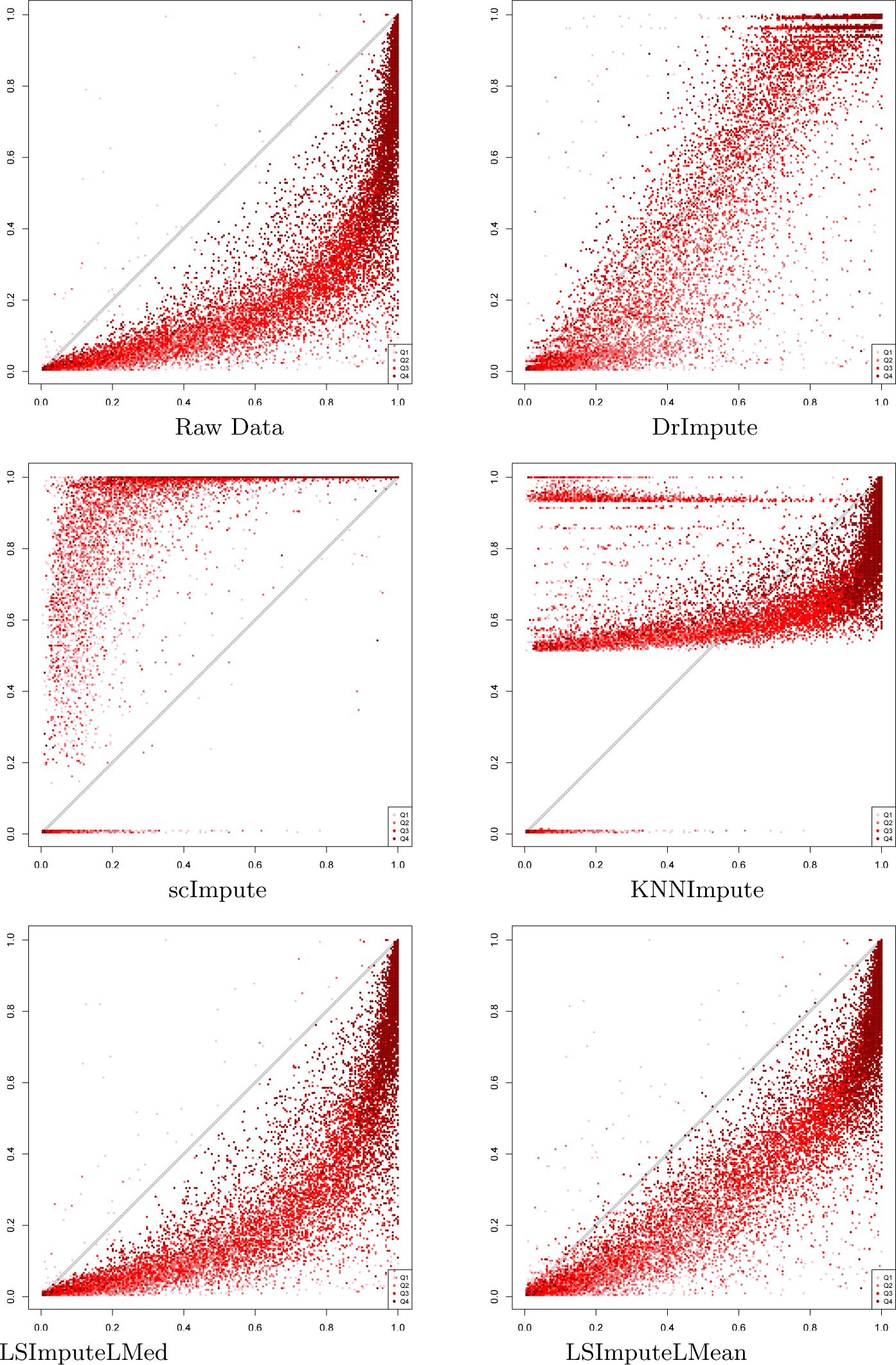
True vs. imputed detection fractions based on rounded TPM values for 50K read pairs per cell.

**Fig. 11.**
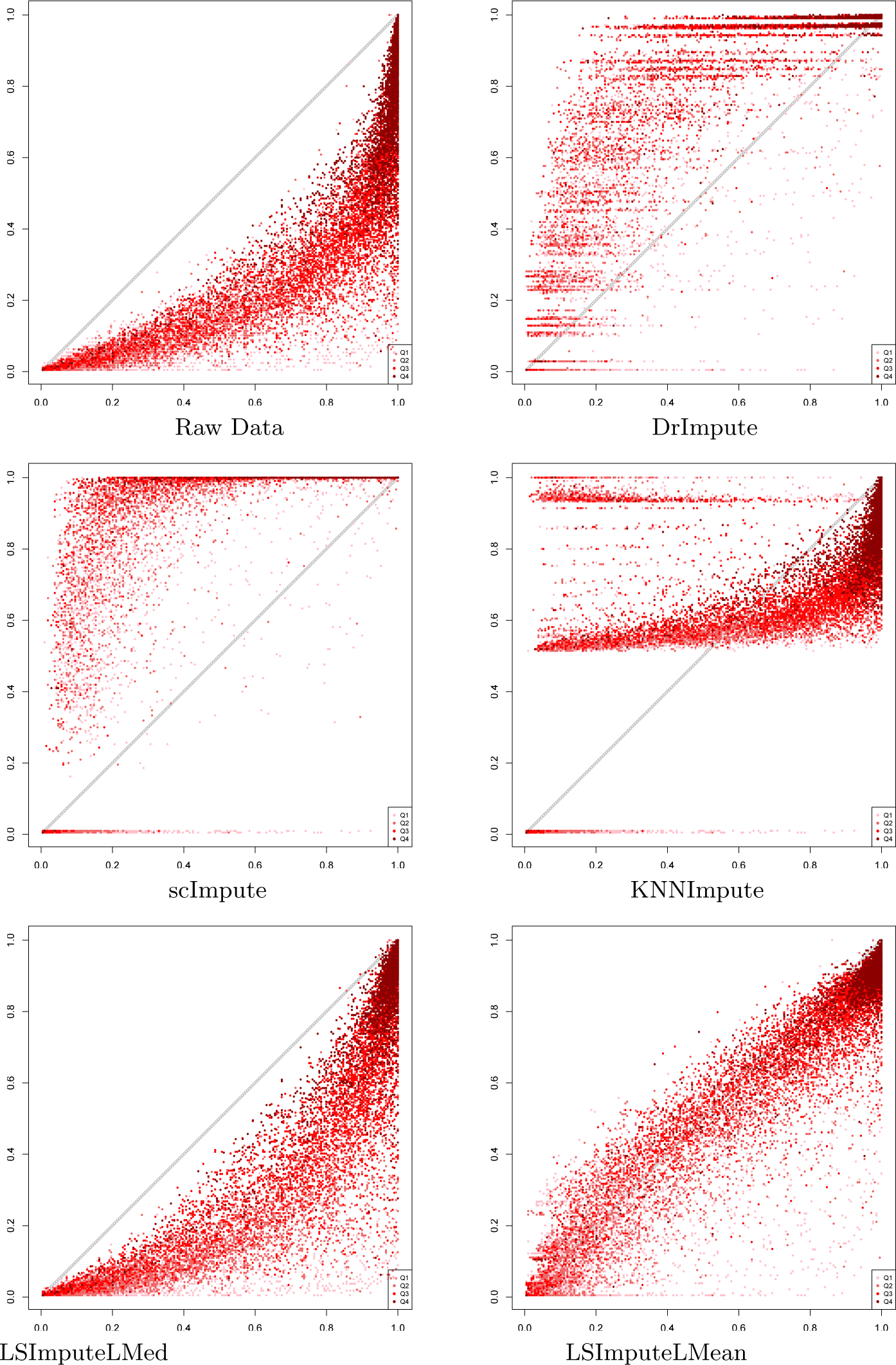
True vs. imputed detection fractions for 100K read pairs per cell.

**Fig. 12.**
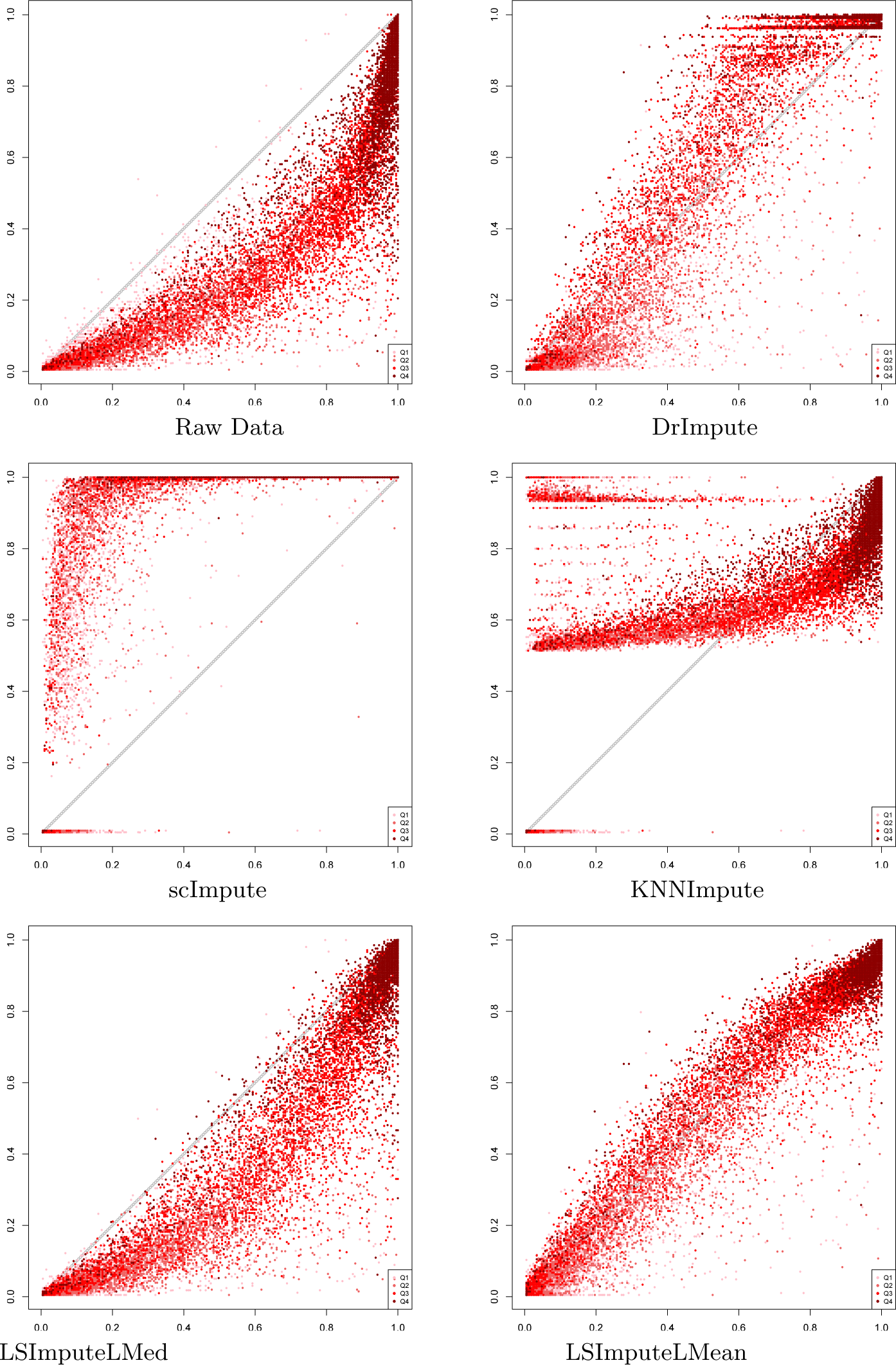
True vs. imputed detection fractions based on rounded TPM values for 100K read pairs per cell.

**Fig. 13.**
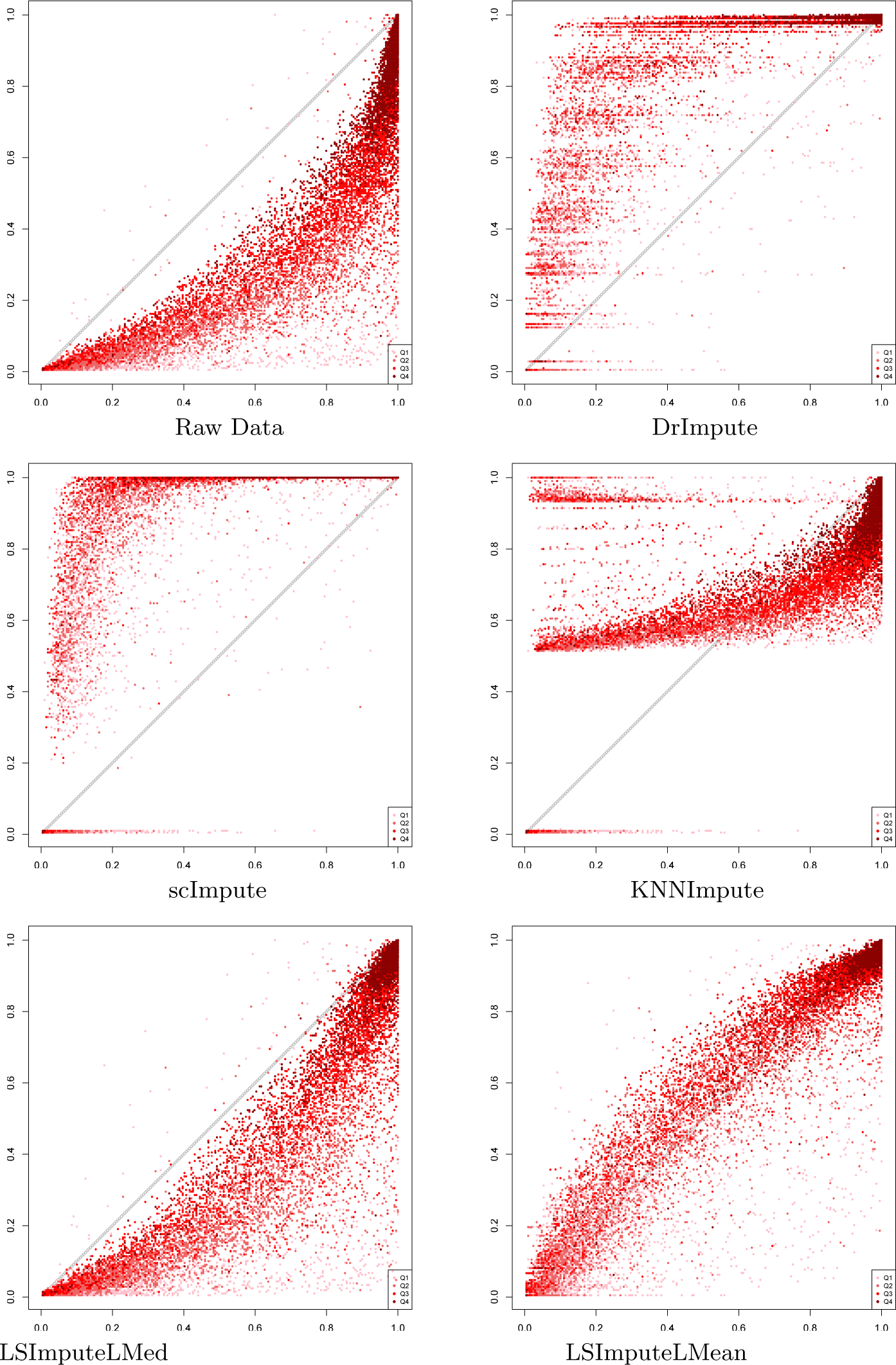
True vs. imputed detection fractions for 200K read pairs per cell.

**Fig. 14.**
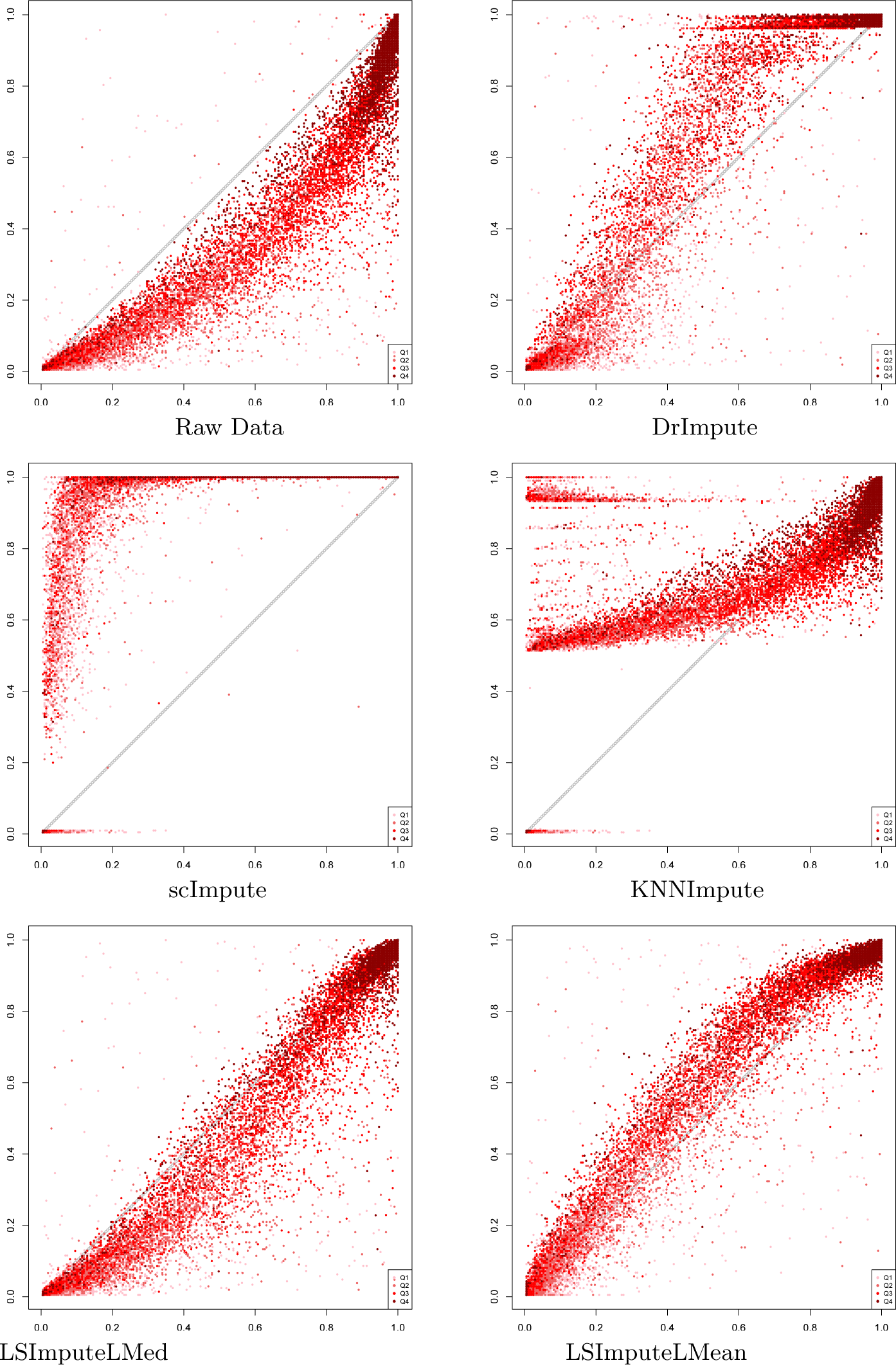
True vs. imputed detection fractions based on rounded TPM values for 200K read pairs per cell.

**Fig. 15.**
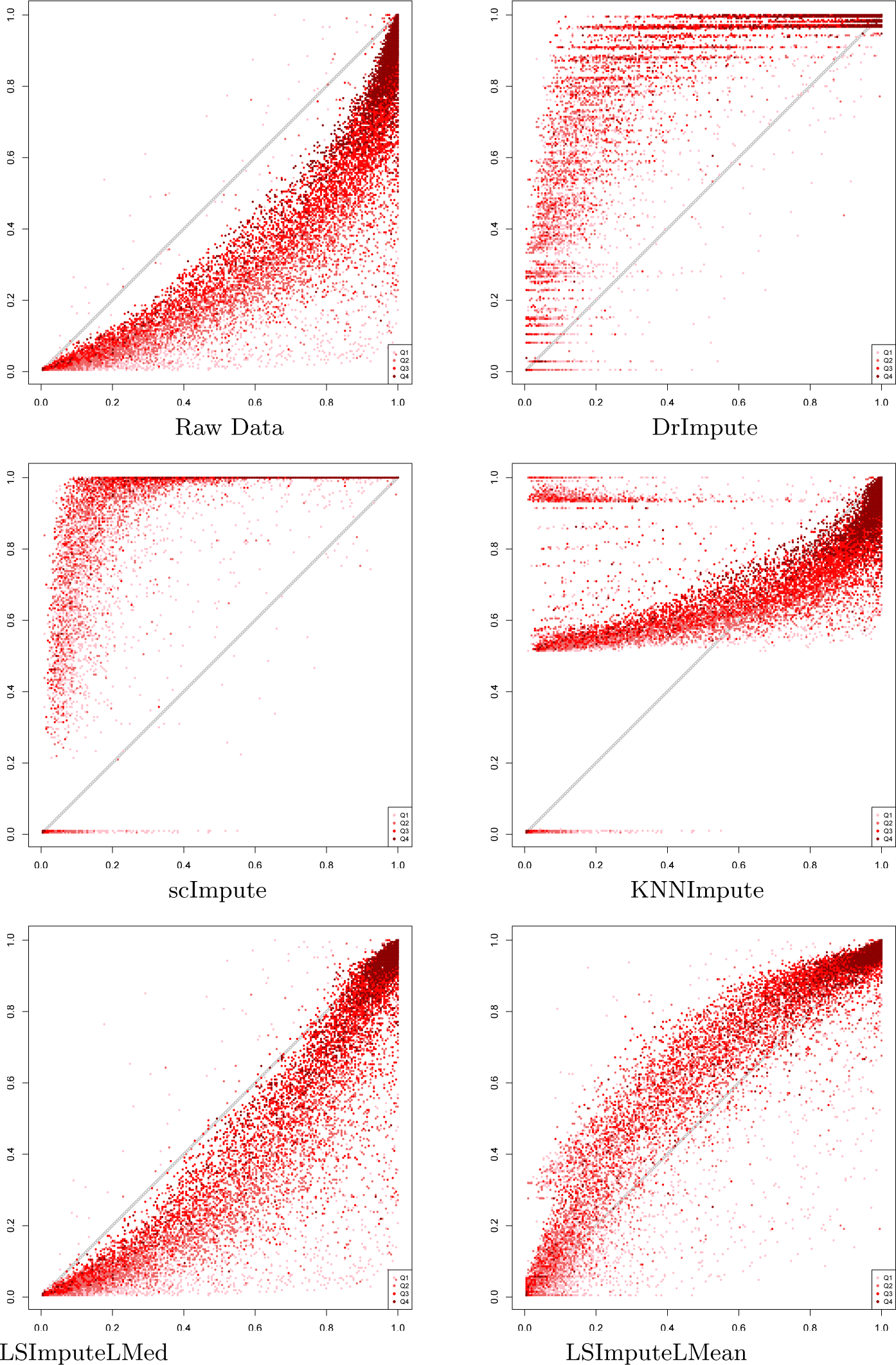
True vs. imputed detection fractions for 300K read pairs per cell.

**Fig. 16.**
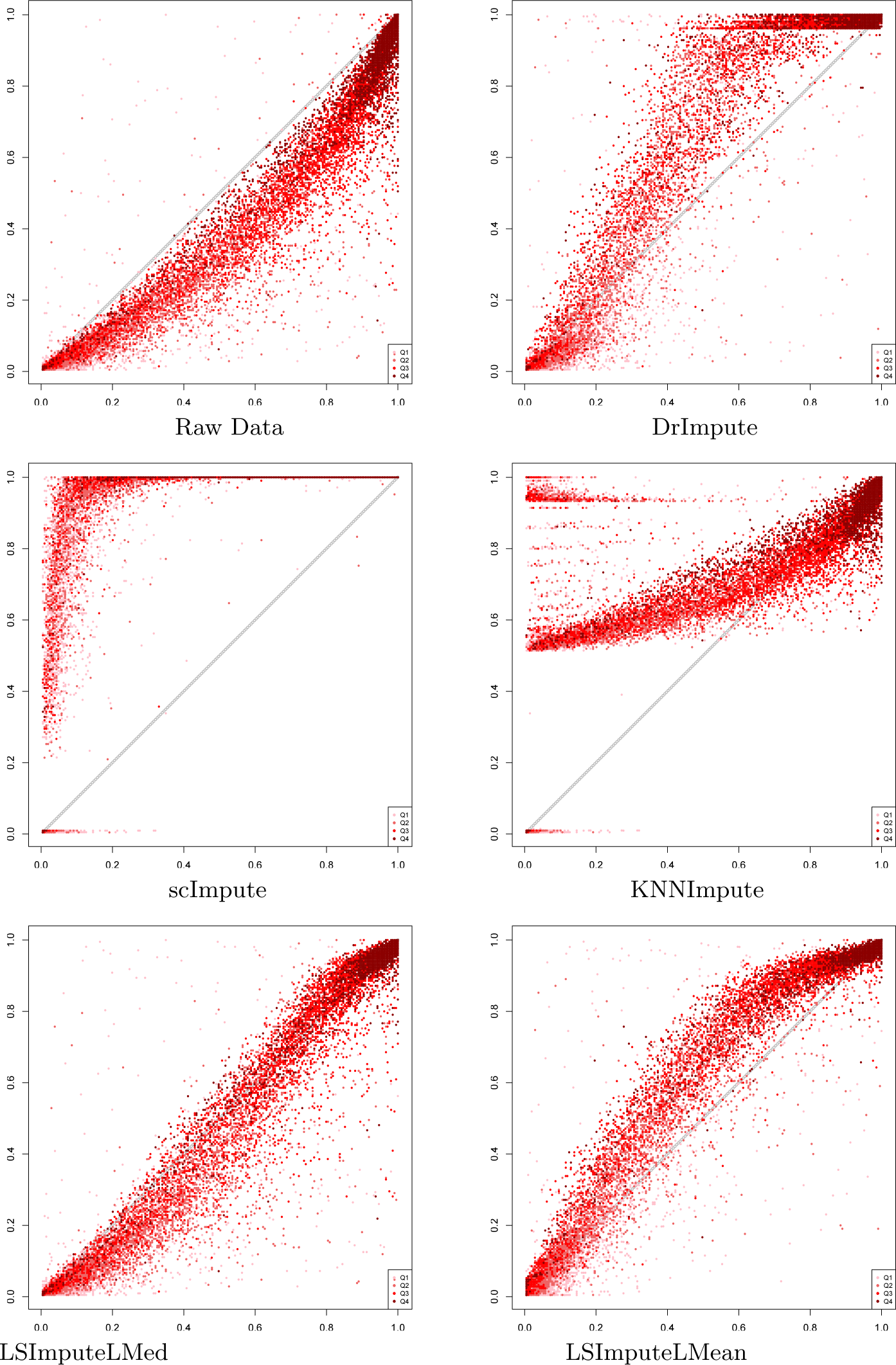
True vs. imputed detection fractions based on rounded TPM values for 300K read pairs per cell.

**Fig. 17.**
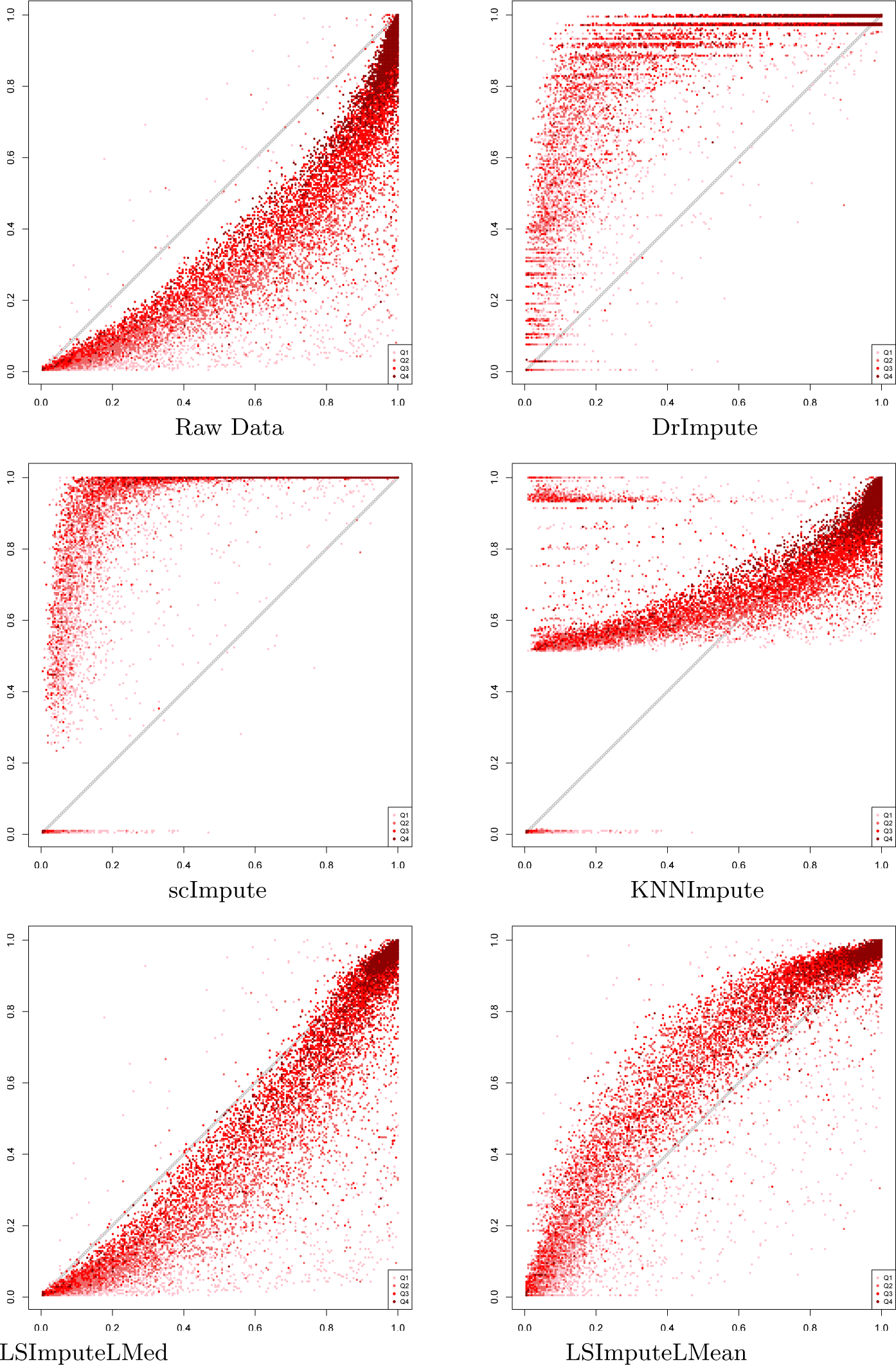
True vs. imputed detection fractions for 400K read pairs per cell.

**Fig. 18.**
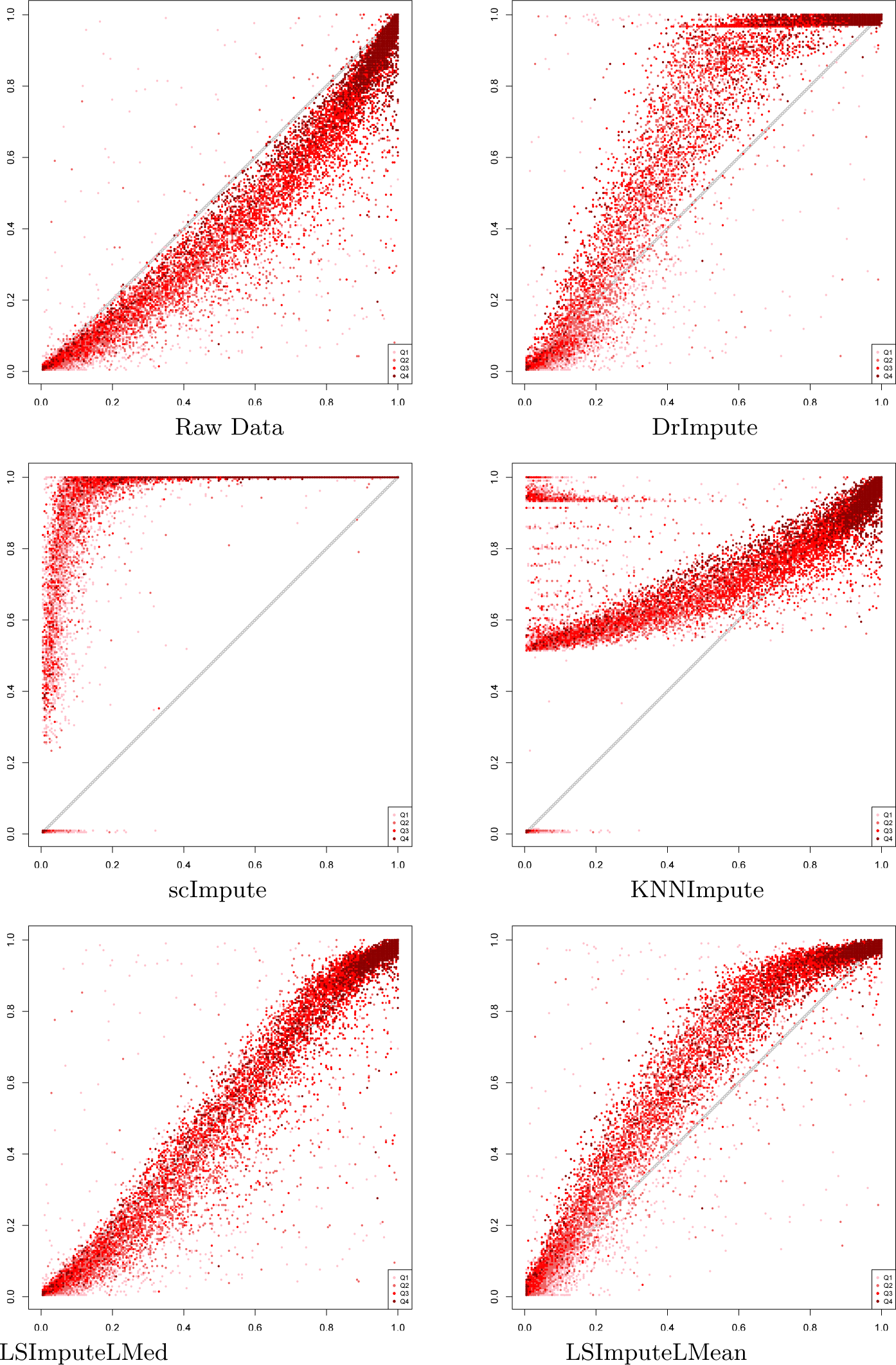
True vs. imputed detection fractions based on rounded TPM values for 400K read pairs per cell.

**Fig. 19.**
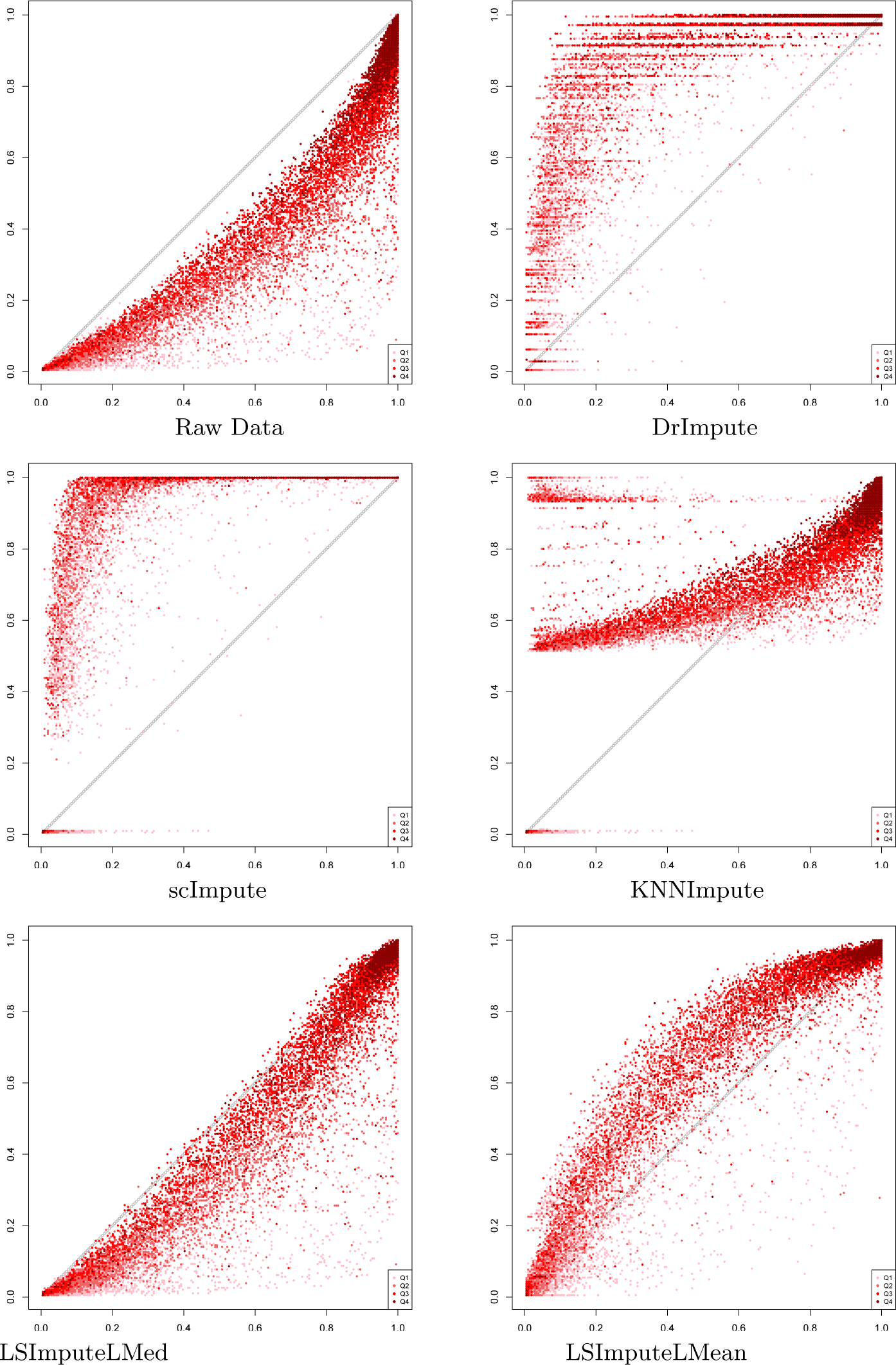
True vs. imputed detection fractions for 500K read pairs per cell.

**Fig. 20.**
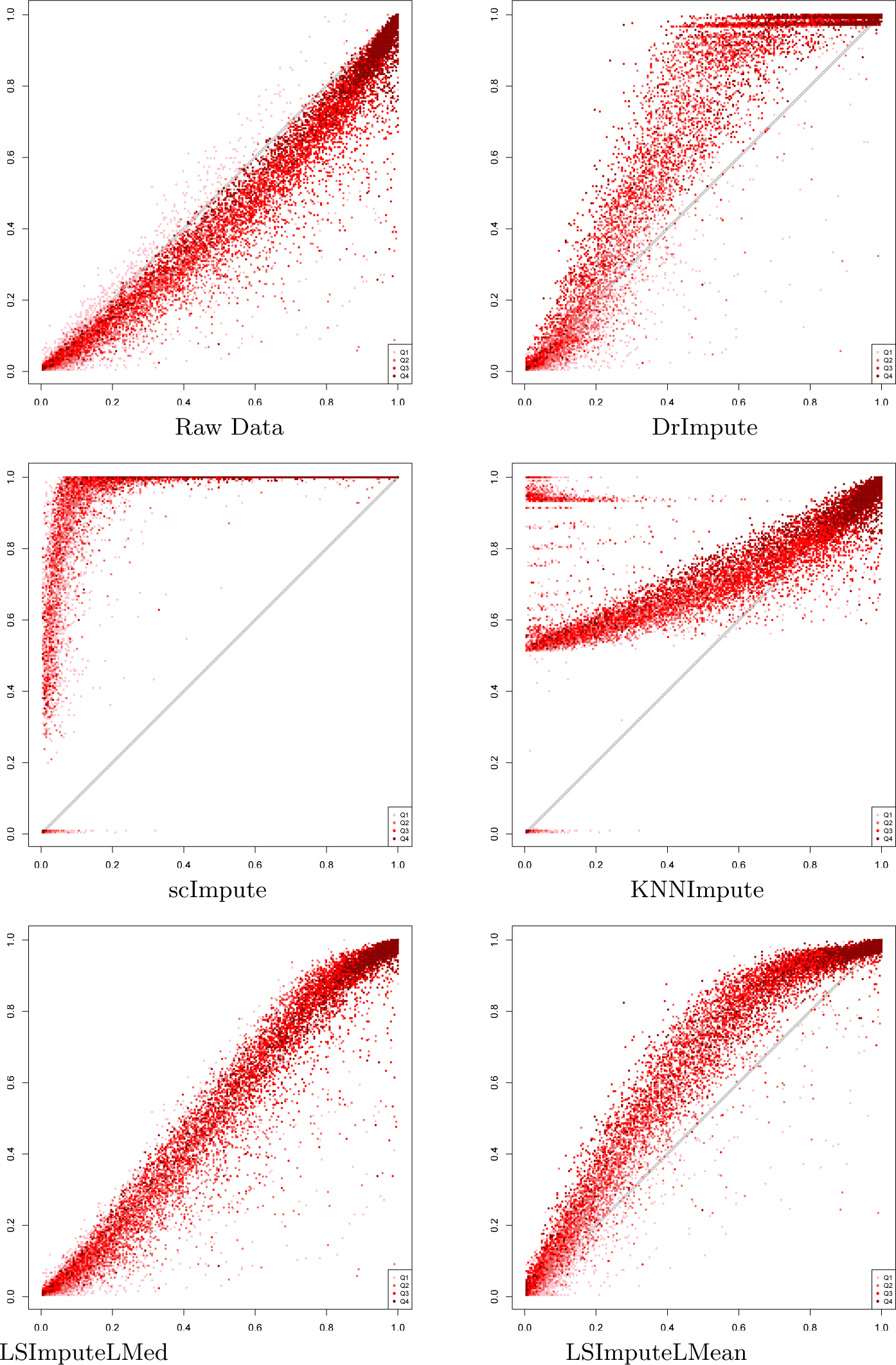
True vs. imputed detection fractions for 500K read pairs per cell.

**Fig. 21.**
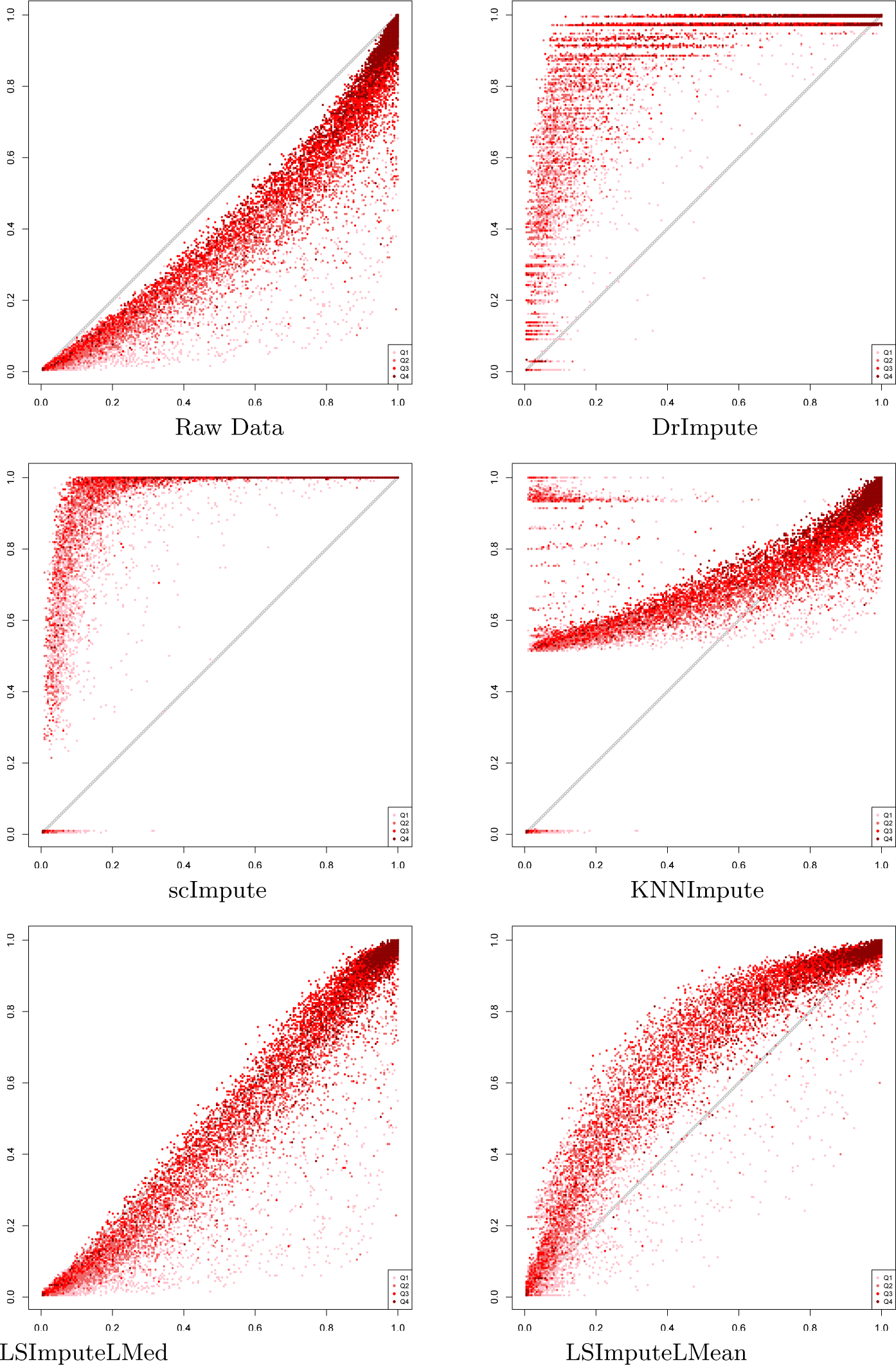
True vs. imputed detection fractions for 1M read pairs per cell.

**Fig. 22.**
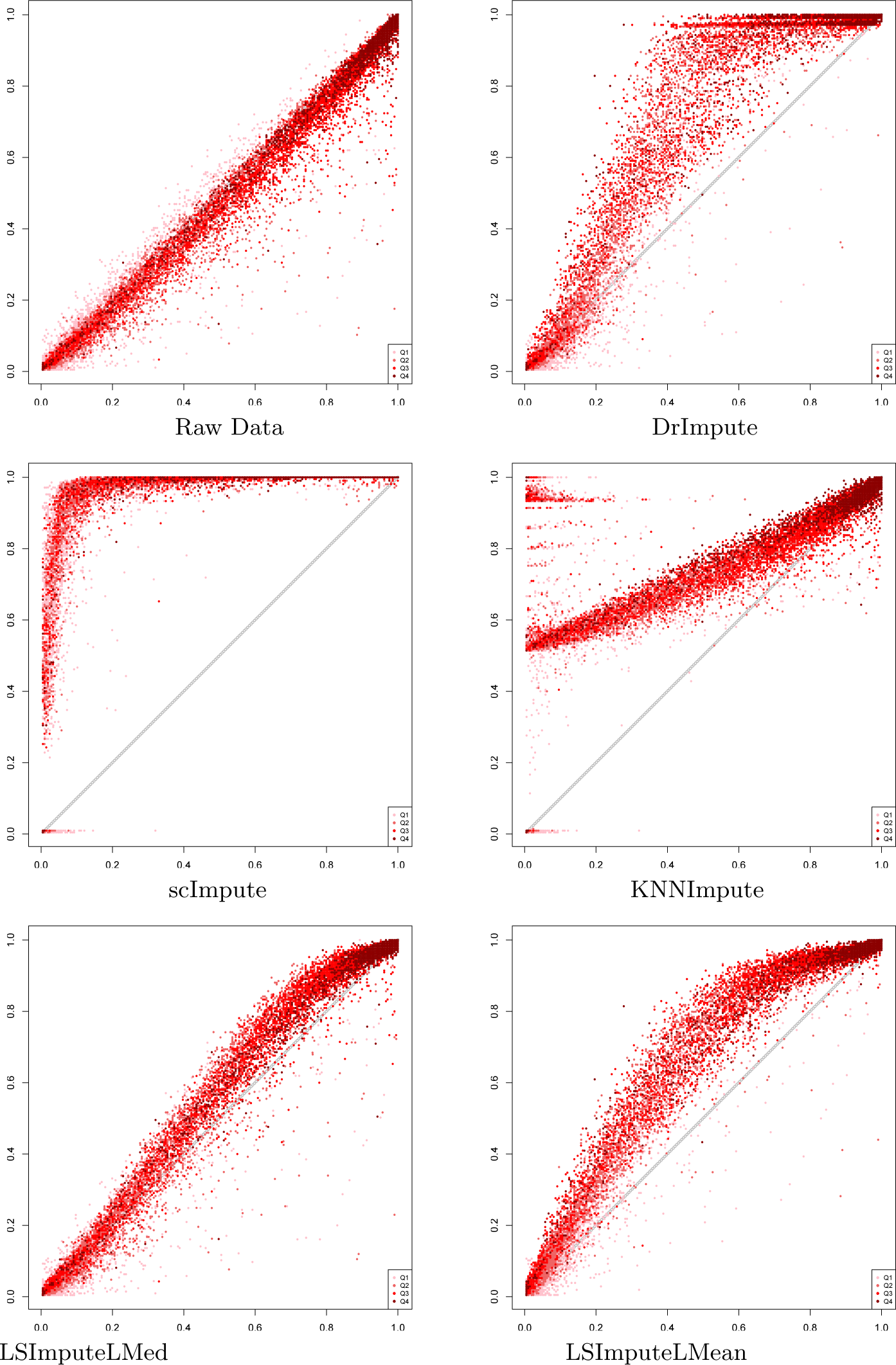
True vs. imputed detection fractions for 1M read pairs per cell.

**Fig. 23.**
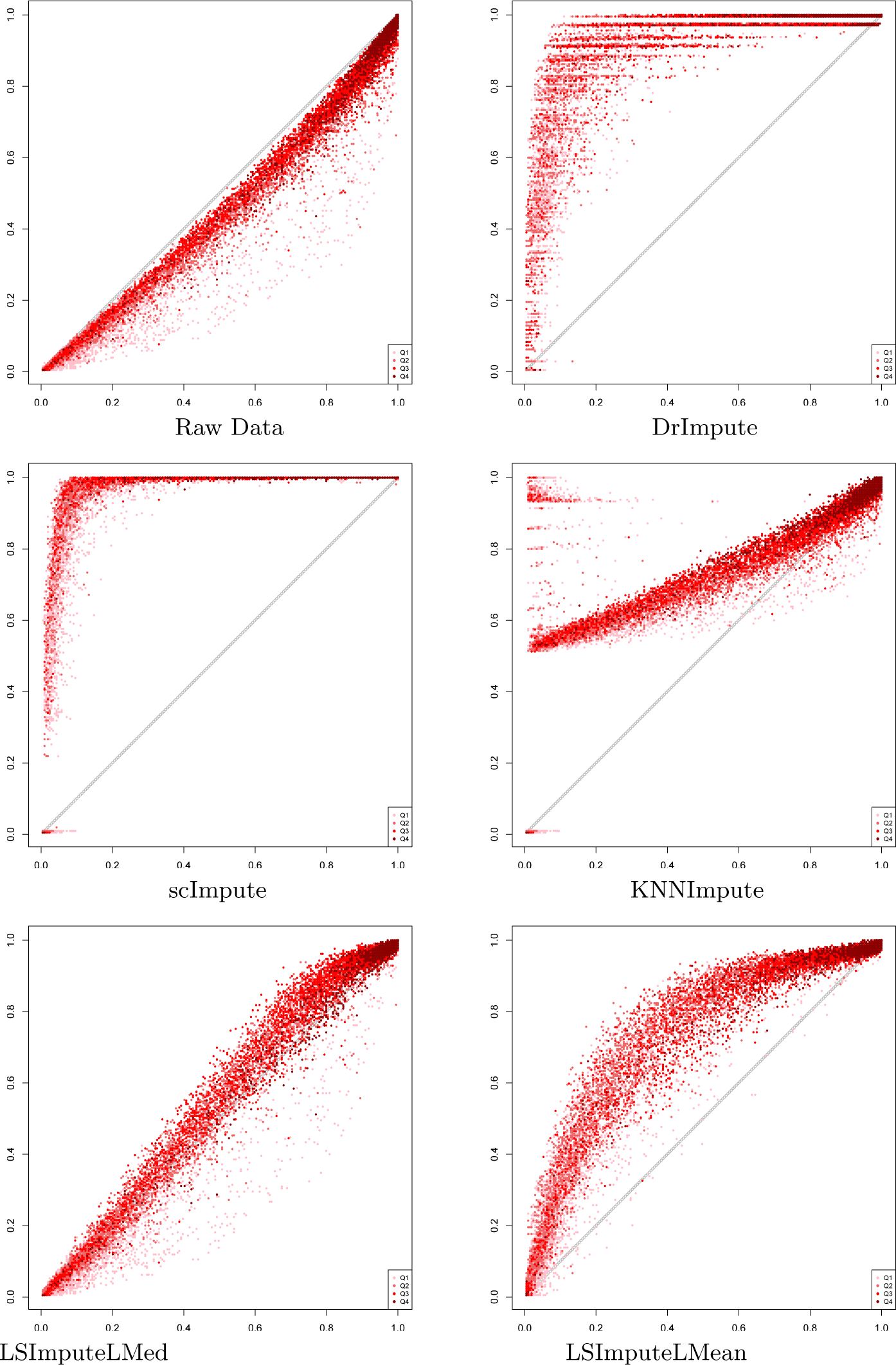
True vs. imputed detection fractions for 5M read pairs per cell.

**Fig. 24.**
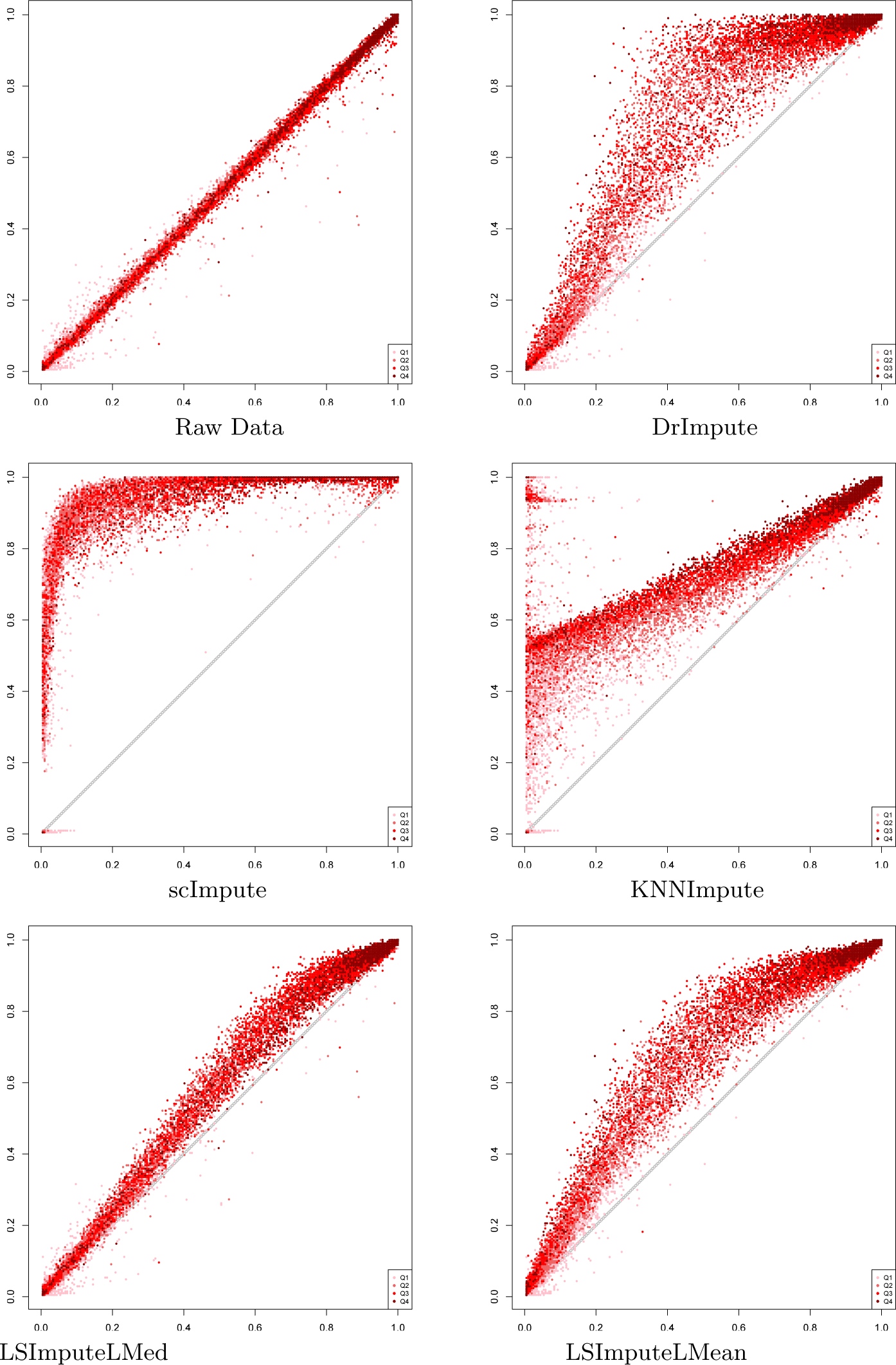
True vs. imputed detection fractions based on rounded TPM values for 5M read pairs per cell.

**Fig. 25.**
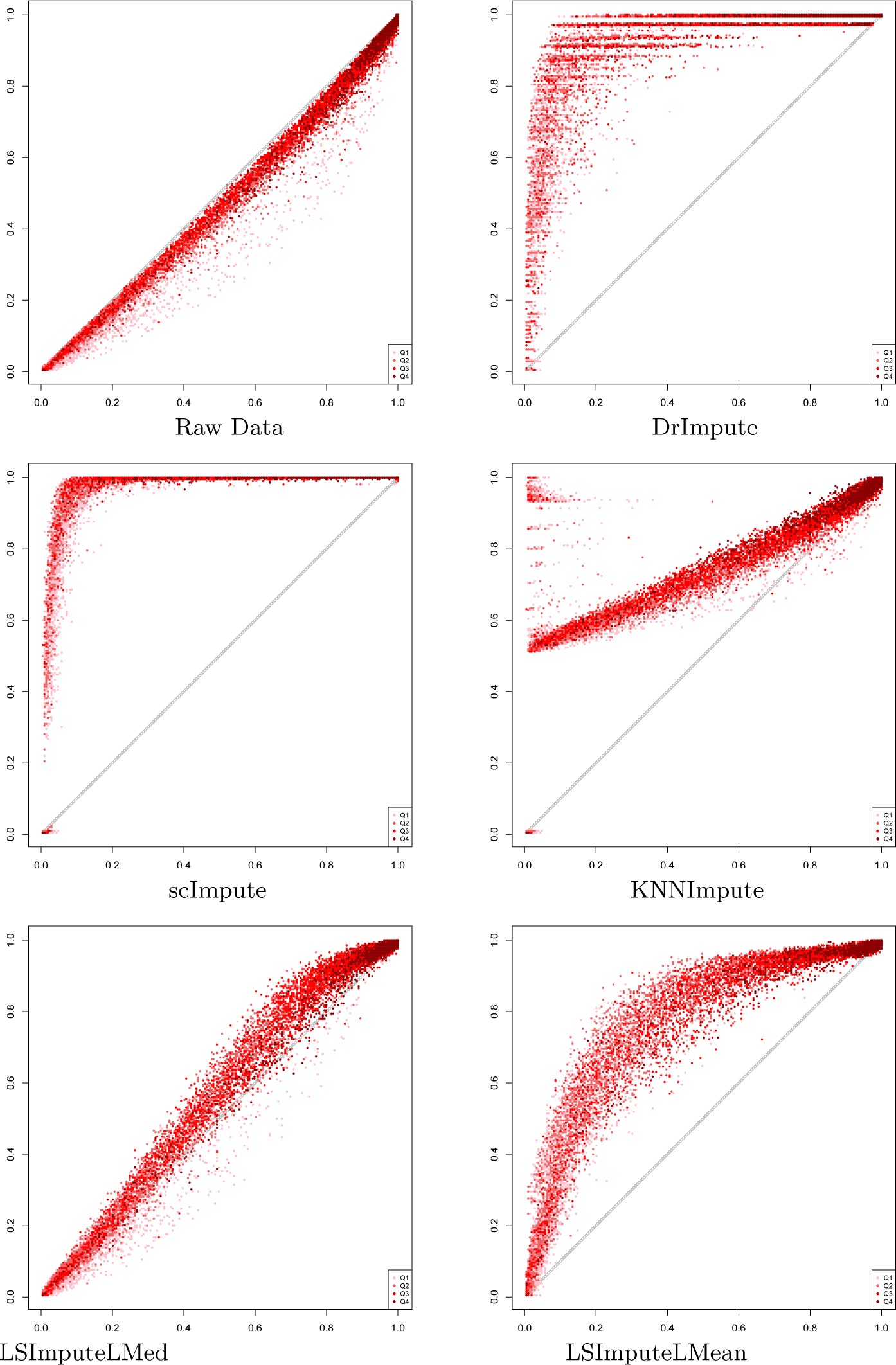
True vs. imputed detection fractions for 10M read pairs per cell.

**Fig. 26.**
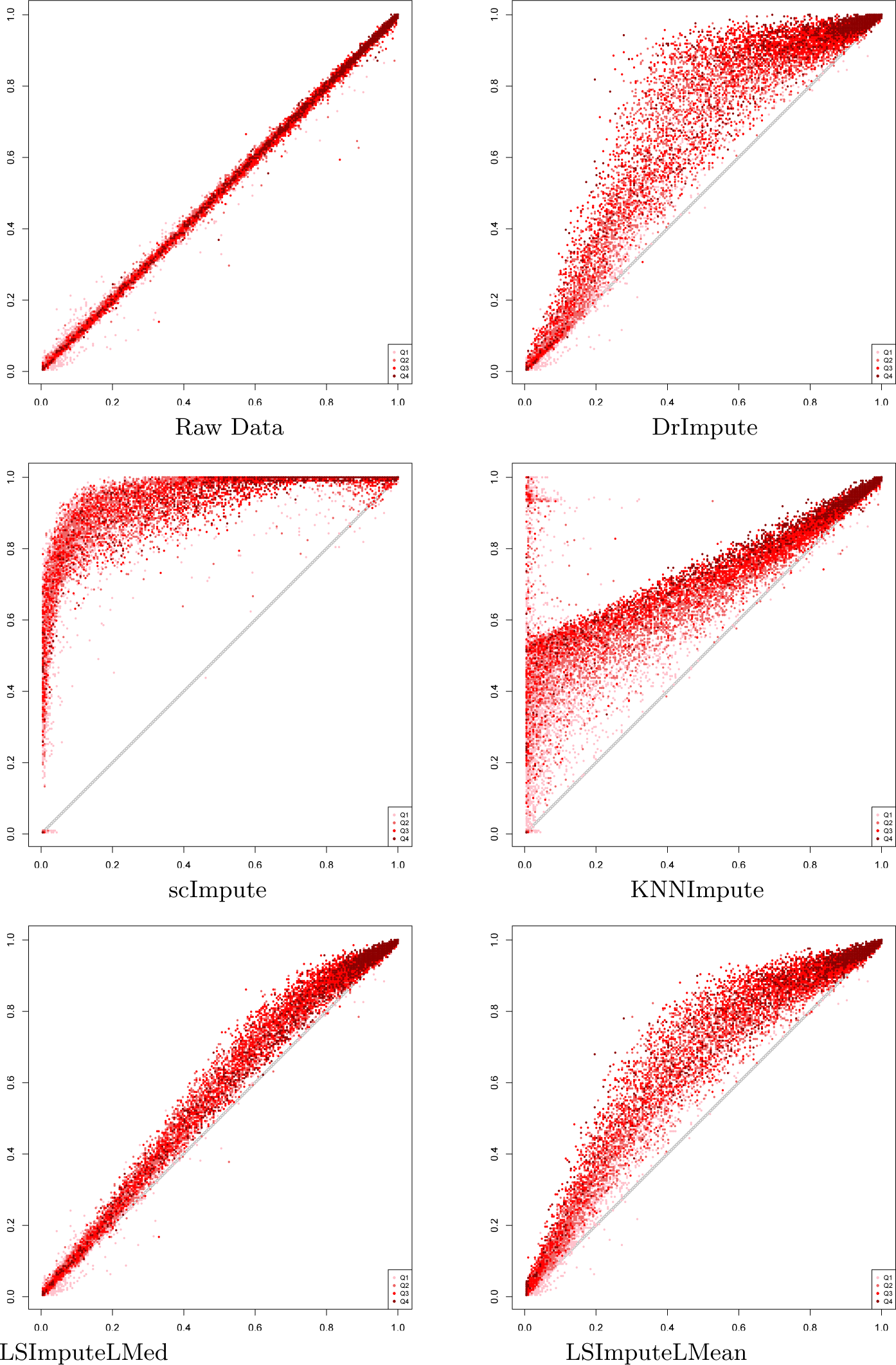
True vs. imputed detection fractions based on rounded TPM values for 10M read pairs per cell.

**Fig. 27.**
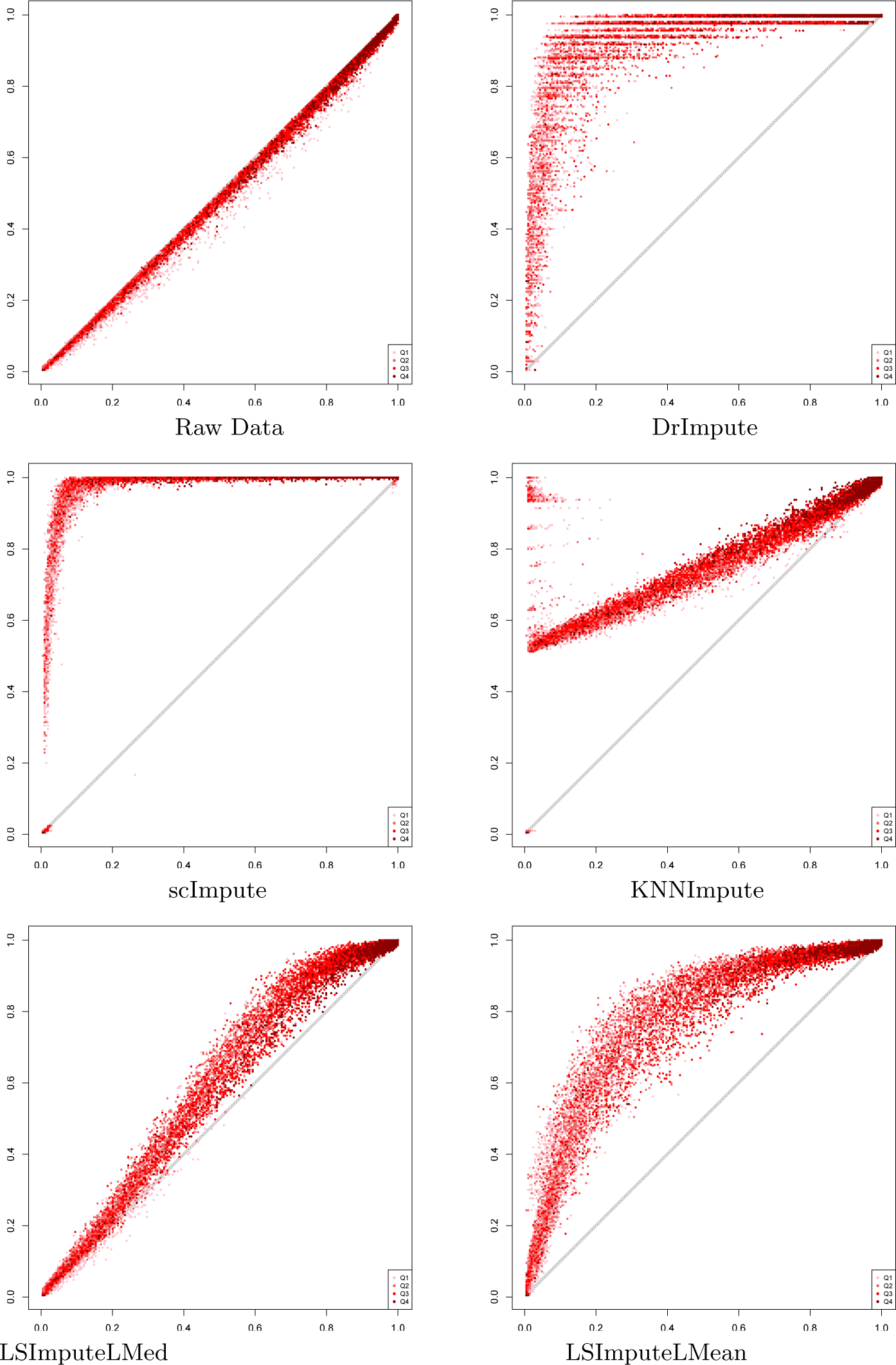
True vs. imputed detection fractions based on rounded TPM values for 20M read pairs per cell.

**Fig. 28.**
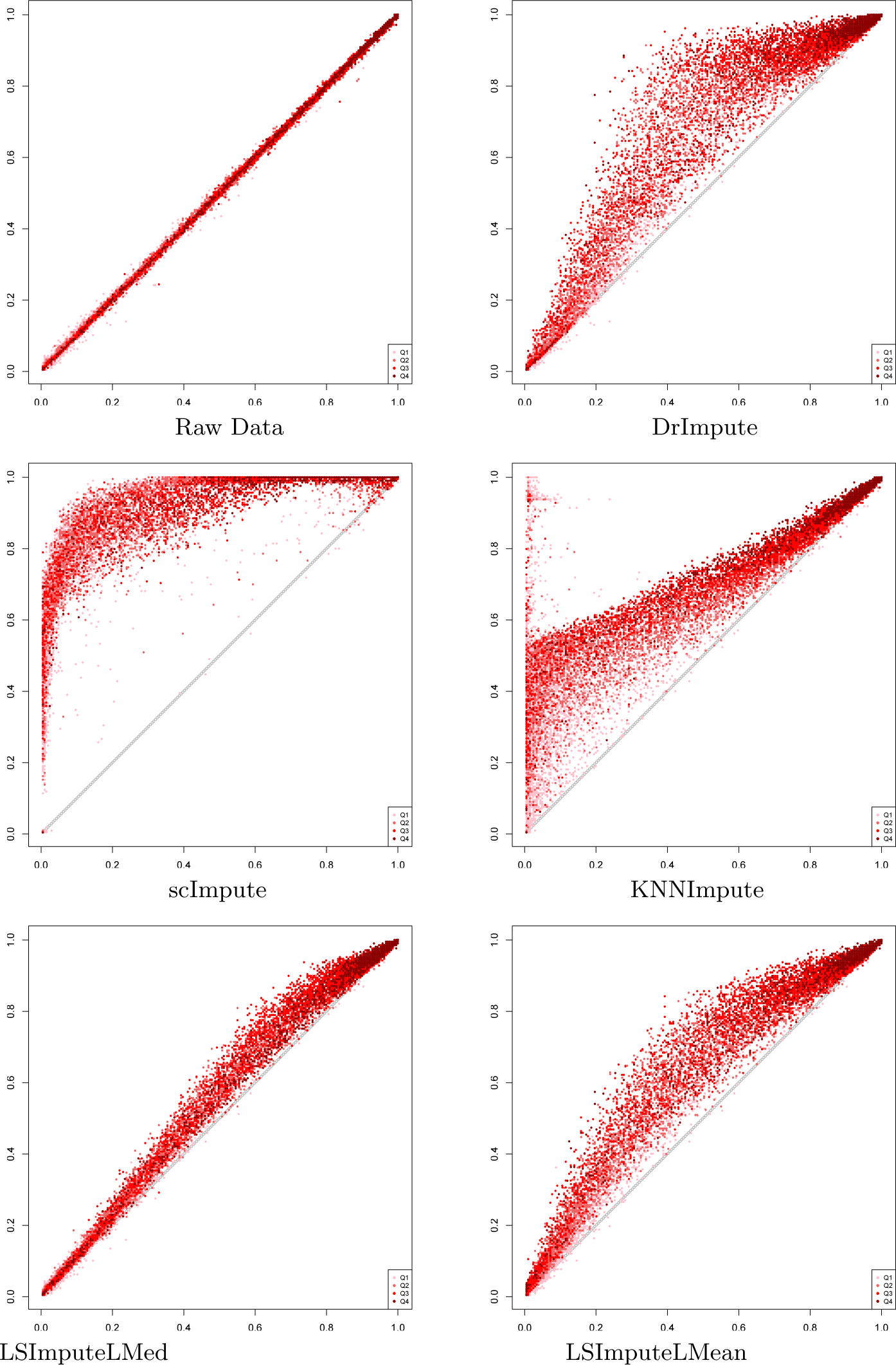
True vs. imputed detection fractions based on rounded TPM values for 20M read pairs per cell.

## Footnotes

1 Note that, unlike KNN, which uses similarity between genes, LSImpute uses similarity between cells. Also, the number of nearest cells used for imputation is not fixed but depends on the minimum similarity threshold.

